# Iron accumulation drives fibrosis, senescence, and the senescence-associated secretory phenotype

**DOI:** 10.1101/2022.07.29.501953

**Authors:** Mate Maus, Vanessa López-Polo, Miguel Lafarga, Mònica Aguilera, Eugenia De Lama, Kathleen Meyer, Anna Manonelles, Anna Sola, Cecilia Lopez Martinez, Ines López-Alonso, Fernanda Hernandez-Gonzales, Selim Chaib, Miguel Rovira, Mayka Sanchez, Rosa Faner, Alvar Agusti, Neus Prats, Guillermo Albaiceta, Josep M. Cruzado, Manuel Serrano

## Abstract

Fibrogenesis is part of a normal protective response to tissue injury that can become irreversible and progressive, leading to fatal diseases. Senescent cells are a main driver of fibrotic diseases through their secretome, known as senescence-associated secretory phenotype (SASP). However, the mechanisms involved in the conversion of damaged cells into senescent cells remain incompletely understood. Here, we report that multiple types of fibrotic diseases in mice and humans are characterized by the accumulation of iron. We show that vascular and hemolytic injuries, through the release of iron, are efficient in triggering senescence and fibrosis. Interestingly, the accumulation of iron is an intrinsic property of senescent cells that does not require an abnormal surge in extracellular iron. Upon damage, cells initiate an iron accumulation response with abundant ferritin-bound iron within lysosomes and high levels of labile iron, the latter being a main driver of senescence-associated ROS and SASP. Finally, we demonstrate that detection of iron by magnetic resonance imaging (MRI) is a powerful non-invasive method to assess fibrotic burden in the kidneys of mice and patients with renal fibrosis. Our findings establish a central role for iron accumulation in senescence and fibrogenesis.

## INTRODUCTION

Fibrosis is the progressive replacement of healthy tissues by a collagen-rich scar tissue^1^. It can affect any organ and manifest in the form of multiple diseases, such as cardiovascular diseases^2^, interstitial lung diseases (ILD)^3^, or kidney disease^4^. Estimates are that fibrotic diseases account for 45% of all mortality in the developed countries^5^. Fibrogenesis is initiated by damage to epithelial and/or endothelial cells, which start secreting chemokines and matrix remodeling enzymes that facilitate the recruitment of macrophages and neutrophils^6^. Neutrophils and macrophages mount an inflammatory response that promotes further matrix remodeling through secretion of matrix metalloproteinases (MMP) and fibrogenic cytokines^7, 8^. A key fibrogenic cytokine is transforming growth factor beta (TGFβ) that is essential for the differentiation and survival of collagen-producing myofibroblasts^9, 10^. While these events are initially reversible, progressive loss of capillarization due to suppression of vascular endothelial growth factor signaling (VEGF) may result in irreparable scar tissue and loss of tissue function^11,12,13^.

Fibrogenesis can be provoked by multiple external insults, such as toxins and infections, but in many fibrotic diseases the etiology remains unknown. Importantly, fibrotic conditions initiated by an external trigger can keep progressing after cessation of the primary insult, as documented in acute kidney injury progressing to chronic kidney disease^14^. This suggests that the primary insult generates pathological entities that persist and are fibrogenic on their own. Multiple reports indicate that senescent cells are abundant in fibrotic tissues and directly contribute to the associated pathologies^15,16,17,18^. Senescent cells are damaged cells characterized by the upregulation of cell cycle inhibitors, more prominently CDKN2A (p16) and CDKN1A (p21), by the presence of high levels of reactive oxygen species (ROS), by a large expansion of the lysosomal compartment, and by their abundant secretion of inflammatory cytokines, chemokines and matrix remodeling enzymes, globally known as senescence-associated secretory phenotype (SASP)^19^. Cellular senescence can limit fibrogenesis when curtailing the proliferation of myofibroblasts^20,21,22^. However, when senescent cells accumulate in tissues they can be fibrogenic on their own^15,16,17,18^ through pro-fibrogenic factors present in the SASP, most notably, TGFβ, IL-11 and SERPINE1^17, 23, 24^.

It is well established that hereditary and acquired hemochromatosis, conditions characterized by the accumulation of iron in tissues, have a high predisposition to develop fibrotic diseases^25,26,27,28,29^. Moreover, iron accumulation has been observed in a number of fibrotic diseases^30,31,32^ and in senescent cells^33, 34^, although the functional role of iron in these processes has not been explored. Here, we propose a unifying connection between cellular damage, iron accumulation, senescence and fibrosis.

## RESULTS

### Tissue injury provokes iron accumulation that progresses throughout fibrosis

We began by assessing whether iron accumulation is a general feature of fibrotic tissues using Perl’s prussian blue staining, either regular or enhanced. We observed iron accumulation in murine lung fibrosis induced by intratracheal bleomycin (**Fig. 1a; and Extended Data Fig. 1a**), and in the lungs of patients with idiopathic pulmonary fibrosis (IPF) (**Fig. 1b**). Similar observations were made in murine kidney fibrosis induced by intraperitoneal folic acid injection (**Fig. 1c**) or unilateral ureteral obstruction (**Extended Data Fig. 1b**), and in kidney biopsies from patients with diabetic kidney disease (**Fig. 1d and Extended Data Table 1**). Iron accumulation was also evident in fibrotic hearts of mice with either transgenic overexpression of the beta adrenergic receptor (*Adrb1*) or ischemic heart disease caused by coronary artery ligation (**Extended Data Fig. 1c and 1d**). Fibrosis-associated iron accumulation was also reflected at the transcriptional level. Bleomycin-induced lung fibrosis and folic acid-induced kidney fibrosis resulted in significantly elevated mRNA levels of the iron-storage protein ferritin (*Fth1* and *Ftl*) (**Extended Data Fig. 1e and 1f**). We also analyzed if there was an enrichment in the n=59 major iron transporter and storage genes (GO: 0006826) in the lung transcriptome of control patients and IPF patients using gene set variation analysis (GSVA). We used two datasets from “The Lung Tissue Research Consortium” (GSE47460)^35^. IPF patients in both datasets showed a marked and significant increase in the iron transport GSVA scores (**Figure 1g**), suggesting that altered gene-expression of iron uptake and storage genes is a defining feature of IPF.

**Fig. 1.**
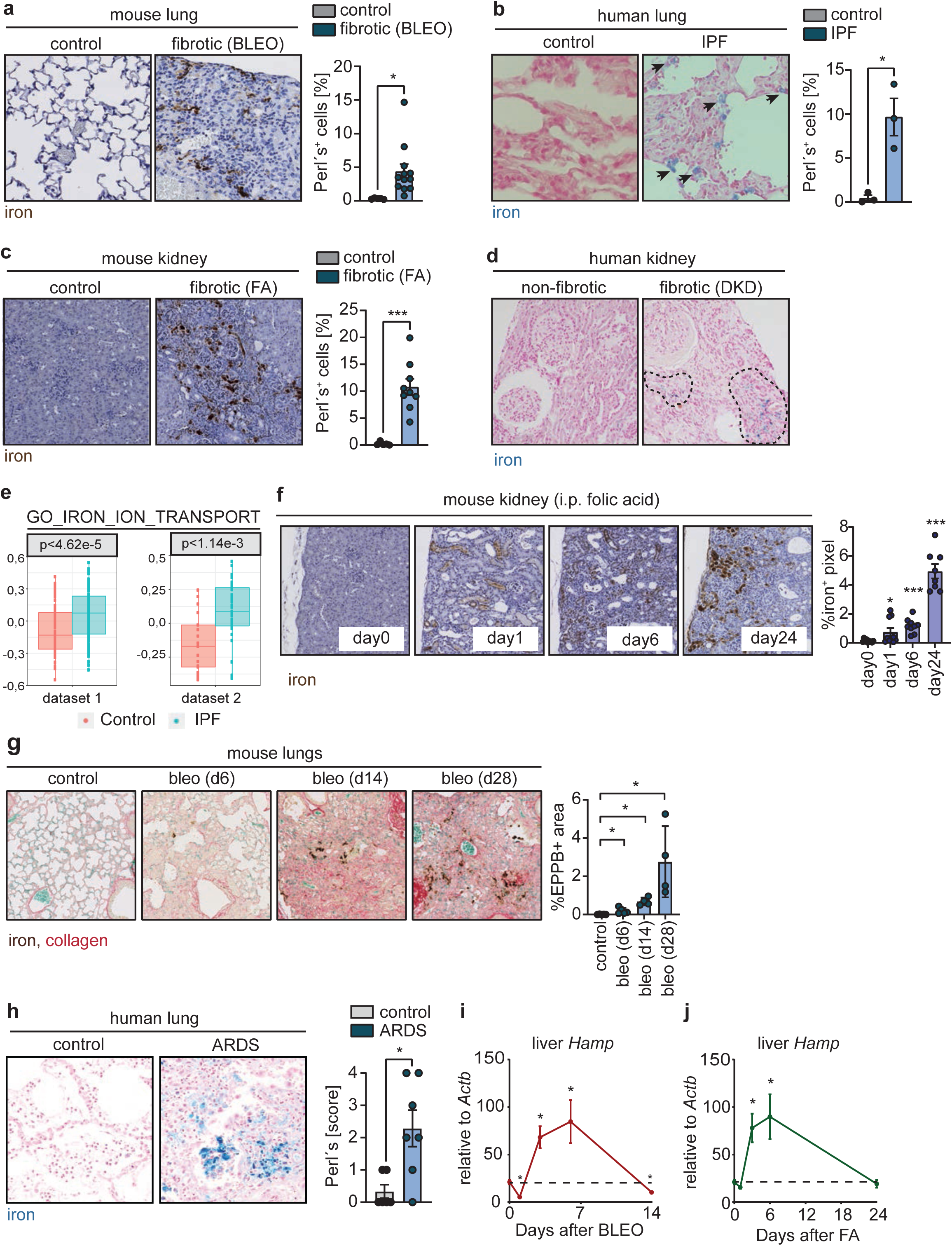
Tissue injury provokes iron accumulation that progresses throughout fibrosis. **a**, Histochemistry for iron by enhanced Perl’s prussian blue staining. Representative images of mouse lung sections 14 days post-intratracheal administration of PBS (control; n=5) or bleomycin (fibrotic; n=12) and quantification. **b**, Histochemistry for iron by regular Perl’s prussian blue staining. Representative images of human lung sections from control patients (no IPF; n=3) and IPF patients (n=3), and quantification. Arrows point at Perĺs positive cells. **c**, Histochemistry for iron by enhanced Perl’s prussian blue staining. Representative images of mouse kidney sections 28 days post-intraperitoneal administration of PBS (control; n=5) or folic acid (fibrotic; n=9) and quantification. **d**, Histochemistry for iron by regular Perl’s prussian blue staining. Representative images of human kidney biopsies from diabetic kidney disease (DKD) patients without (non- fibrotic) or with interstitial fibrosis (fibrotic). Dashed lines delineate the fibrotic area abundant in Perĺs positive cells. Quantification in Table 1. **e**, Boxplots of the Gene-set variation analysis (GSVA) relative expression scores of the iron transport and storage pathway (GO_IRON_ION_TRANSPORT:0006826; n=59 genes) computed on two different whole lung transcriptome datasets from controls (dataset1, n=78; dataset2, n=17) and IPF patients (dataset1, n=122; dataset2, n=38). **f**, Histochemistry for iron by enhanced Perl’s prussian blue staining. Representative images of mouse kidneys from before (n=7), and 1 day (n=9), 6 days (n=9) or 24 days (n=7) post-intraperitoneal administration of folic acid, and quantification. **g**, Histochemistry for iron and collagen by combining enhanced Perl’s prussian blue and Sirius Red/Fast Green staining. Representative images of mouse lung sections from before (n=5), and 6 days (n=5), 14 days (n=4) or 28 days (n=4) post-intratracheal administration of bleomycin, and quantification. **h**, Histochemistry for iron by regular Perl’s prussian blue staining. Representative images of human lung sections collected post-mortem from control patients (no ARDS; n=6) and ARDS patients (n=7), and quantification. **i-j**, mRNA levels of hepcidin (*Hamp*) relative to *Actb* in the livers of mice **i**, from before (n=6), or 1 day (n=5), 3 days (n=5), 6 days (n=4), or 14 days (n=6) post-bleomycin or **j**, from before (n=6), and 1 day (n=5), 3 days (n=5), 6 days (n=5), or 24 days (n=6) post-folic acid. Bar graphs in each case represent mean ± s.e.m.; individual values for each mice or humans are represented as points; in **e** we used the kruskal-wallis test; in **h** we used the Wilcoxon signed- rank test; in all other panels, we used two-tailed unpaired t-test. *p < 0.05; **p < 0.01; ***p < 0.001; ****p < 0.0001.

We assessed the kinetics of iron accumulation throughout fibrogenesis. Iron deposition visualized by enhanced Perĺs Prussian blue staining was detectable in the kidneys of folic acid-treated mice as early as one day after the fibrogenic insult and kept on progressing until the last day of analysis (**Fig. 1f**). We made similar observations in the lungs of bleomycin-treated mice (**Fig. 1g**). We also investigated tissues from patients with acute respiratory distress syndrome (ARDS), a condition that can be caused by multiple independent insults and that often develops into established lung fibrosis^36^. Lungs were analyzed within a few days after ARDS, before development of fibrosis and, interestingly, iron accumulation was observed in most of them irrespective of the type of lung injury that caused ARDS (septic shock in association with pneumonia or kidney transplantation, hemorrhagic shock in association with aortic aneurysm, acute leukemia with systemic mycosis, pneumonia in association with Influenza A, or unknown origin)^37^ (**Fig. 1h, Extended Data Table 2**).

We wondered if the progressive accumulation of iron in damaged tissues had a correlate detectable systemically. Mice treated with intratracheal bleomycin or intraperitoneal folic acid presented a rapid but transient peak of hepatic hepcidin (*Hamp*, measured as mRNA) (**Fig 1i, j**), a hormone that is induced in the liver when iron is in excess^38^. Our findings suggest that iron accumulation is initiated soon after tissue injury and progresses throughout fibrosis. Of note, iron accumulation in the fibrotic lesions progresses over time, while the systemically detectable response to iron excess is transient.

### Iron initiates fibrogenesis

We wondered whether iron accumulation is a bystander phenomenon, or plays a causative role in fibrogenesis. Intratracheal delivery of iron into mice provoked a robust pro-fibrotic cytokine response within two days after injury, as measured by a multiplex protein detection assay (**Fig. 2a**). This response was dominated by G-CSF, a recruitment factor for fibroblasts and granulocytes^39^; members of the pro-fibrotic IL-6 family^40, 41^, including IL-6, IL-11, and LIF; TIMP-1, a collagenase inhibitor that prevents degradation of newly formed collagen^42^; PAI-1 (SERPINE1), a master regulator of collagen deposition in fibrotic diseases^43^; the eosinophil chemoattractant IL- 5^44^; macrophage chemoattractants CCL2 and CCL12; and the neutrophil chemoattractant CXCL1^1, 45, 46^ (**Fig. 2a**). Accordingly, we observed a massive influx of macrophages (**Fig. 2b and Extended Data Fig. 1a**) and neutrophils (**Fig. 2b and Extended Data Fig. 1b**) in iron-rich foci. Interestingly, iron also induced depletion of CD31^+^ endothelial cells detectable by flow-cytometry (**Fig. 2c**) and mRNA levels (**Extended Data Fig. 2c**). Histological analyses showed that loss of CD31^+^ cells mainly affected inflamed foci with immune infiltration (**Fig. 2d**). Depletion of endothelial cells was concomitant with a robust and persistent suppression of VEGF protein and mRNA levels (**Fig. 2a, Extended Data Fig. 2d**), reminiscent of fibrosis-associated vascular rarefaction^11,12,13^. Iron also provoked the remodeling of the extracellular matrix, first visible within two days through the increased abundance in TIMP-1 (**Fig. 2a**) and matrix metalloproteinases (MMP8, 9 and 3) (**Extended Data Fig. 2e**). This was followed by the upregulation of protein levels of the profibrogenic TGFβ family members within 6 days of iron delivery (**Fig. 2e**). At this stage, iron rich foci in the lungs presented with a marked expansion of α-SMA positive myofibroblasts, associated with robust accumulation of collagen (**Fig. 2f**). Two weeks after iron delivery, *Col1a1* and *Col1a2* (collagens) and *Fn1* (fibronectin) mRNA levels were significantly elevated (**Extended Data Fig. 2f**), but histologies showed signs of partial repair, as indicated by a reduction in the size of iron and collagen-rich fibrotic foci (**Extended Data Fig. 2g**). Despite partial repair, iron-rich lesions associated with fibrosis persisted even 28 days after iron delivery (**Extended Data Fig. 2g**). Considering the strong association between fibrosis and cellular senescence^15^, we also tested whether intratracheal iron could induce senescence. Iron triggered a remarkable accumulation of p21^+^ cells in the fibrotic lesions 6 days post-delivery (**Fig. 2g**). Two weeks after iron delivery, we also observed a significant increase in the mRNA levels of *Cdkn2a* (p16) in the lungs of mice treated with iron (**Fig. 2h**). Together, we conclude that iron accumulation is sufficient to initiate hallmarks of fibrogenesis, inflammation, innate immune infiltration, vascular rarefaction, senescence and collagen deposition.

**Fig. 2.**
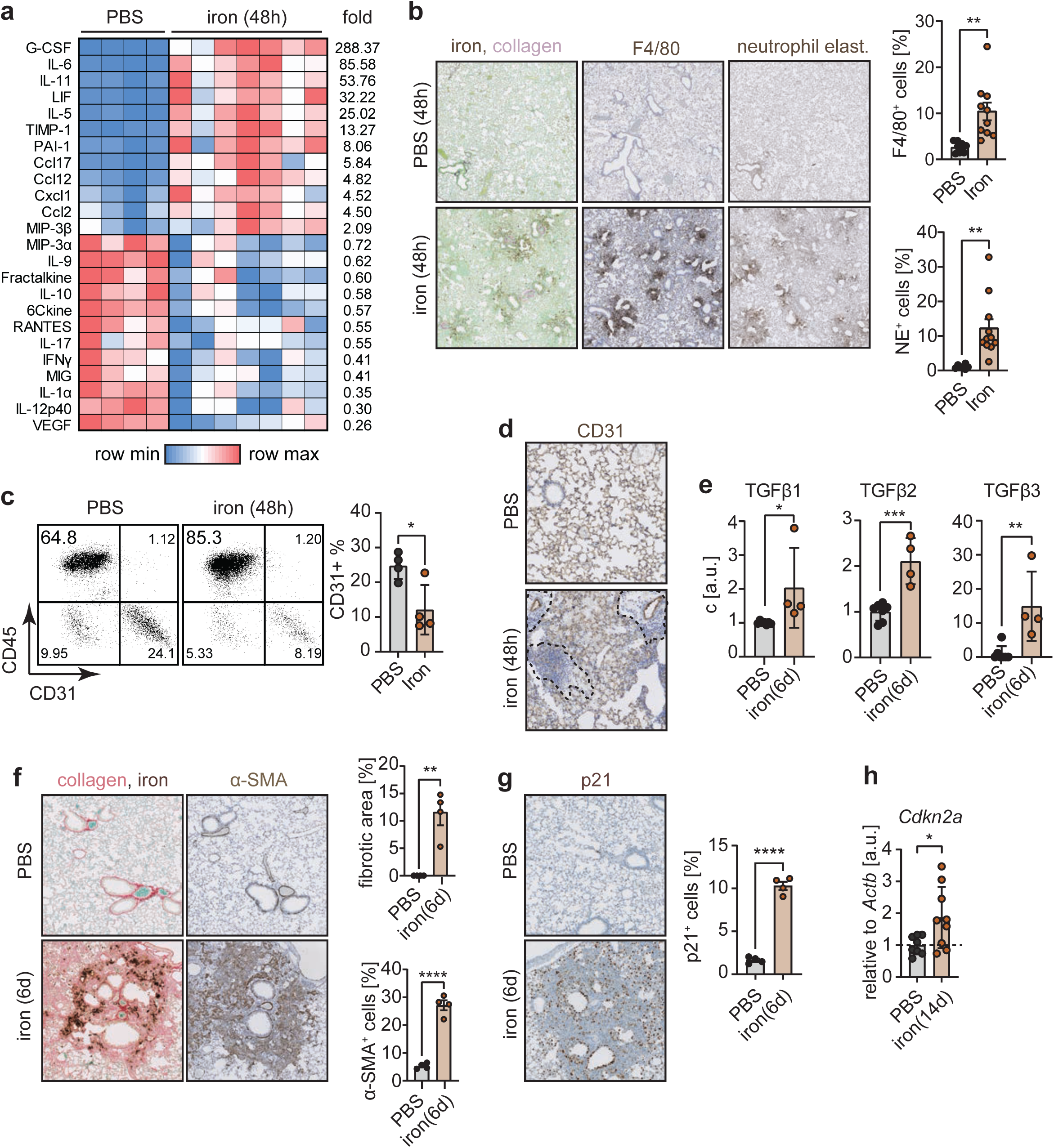
Iron initiates fibrogenesis. **a,** Heatmap of significantly altered cytokines and chemokines in the lungs of mice 2 days post-intratracheal delivery of PBS (n=4) or iron (500 nM; n=7). Lung lysates were tested by a multiplex protein assay for 45 cytokines out of which 24 were significantly changed, with a fold-change as shown to the right. **b,** Histochemistry for iron and collagen, by combining enhanced Perl’s prussian blue and Sirius Red/Fast Green staining, and immunohistochemistry for the macrophage marker F4/80, and for the neutrophil marker, neutrophil elastase. Representative images of mouse lung sections 2 days post-intratracheal administration of PBS (n=6-9) or iron (500 nM; n=9-11). **c**, Flow-cytometric analysis of dissociated lung tissues for the markers CD31 and CD45. Representative dot plots of lung cells from mice 2 days post-intratracheal delivery of PBS (n=4) or iron (500 nM; n=4), and quantification. **d**, Representative images showing immunohistochemical analysis of CD31+ endothelial cells in lung sections from mice 2 days post post-intratracheal delivery of PBS (n=4) or iron (500 nM). Areas surrounded by dashed lines indicate regions where CD31 staining gets lost, which coincides with areas of immune-infiltration. **e**, Levels of TGFβ family members in the lungs of mice 6 days after intratracheal delivery of PBS (n=6-7) or iron (500 nM; n=4). Lung lysates were tested by a multiplex protein assay for the levels of the three TGFβ family members. **f**, Histochemistry for iron and collagen, by combining enhanced Perl’s prussian blue and Sirius Red/Fast Green staining, and immunohistochemistry for the myofibroblast marker α-SMA. Representative images of mouse lung sections 6 days post-intratracheal administration of PBS (n=4) or iron (500 nM; n=4), and quantification. **g**, Immunohistochemistry for the senescence marker p21. Representative images of mouse lung sections 6 days post-intratracheal administration of PBS (n=4) or iron (500 nM; n=4), and quantification. **h**, mRNA levels of *Cdkn2a* (p16) relative to *Actb* in mouse lungs 14 days post-intratracheal administration of PBS (control; n=8) or iron (500 nM; n=9). Bar graphs represent mean ± s.e.m.; individual values for each mouse are represented as points; we used two-tailed unpaired t-test. *p < 0.05; **p < 0.01; ***p < 0.001; ****p < 0.0001.

### Vascular and hemolytic insults provoke iron accumulation, senescence and fibrogenesis

Considering the capacity of iron to induce fibrosis, we asked if vascular or hemolytic injuries could cause iron release from damaged red blood cells (RBC), and thereby inflammation, senescence and fibrosis. To directly interrogate whether vascular injuries are sufficient to drive iron accumulation and senescence *in vivo*, we generated a mouse model (*Tie2-ERT2/CRE; Rosa26- DTR-STOP^fl/fl^*) in which a fraction of endothelial cells can be ablated by the injection of tamoxifen and diphtheria toxin leading to microhemorrhages, mainly in the lungs, compatible with mouse survival. Sites of microvascular injuries were visible as regions with abnormal CD31 staining and extravascular Ter119^+^ RBCs (**Fig. 3a and Extended Data Fig. 3a**). Interestingly, these regions contained abundant iron-loaded cells and p21^+^ cells (**Fig. 3a and Extended Data Fig. 3a**), suggesting a causative role of vascular injury in iron accumulation and senescence *in vivo*.

**Fig. 3.**
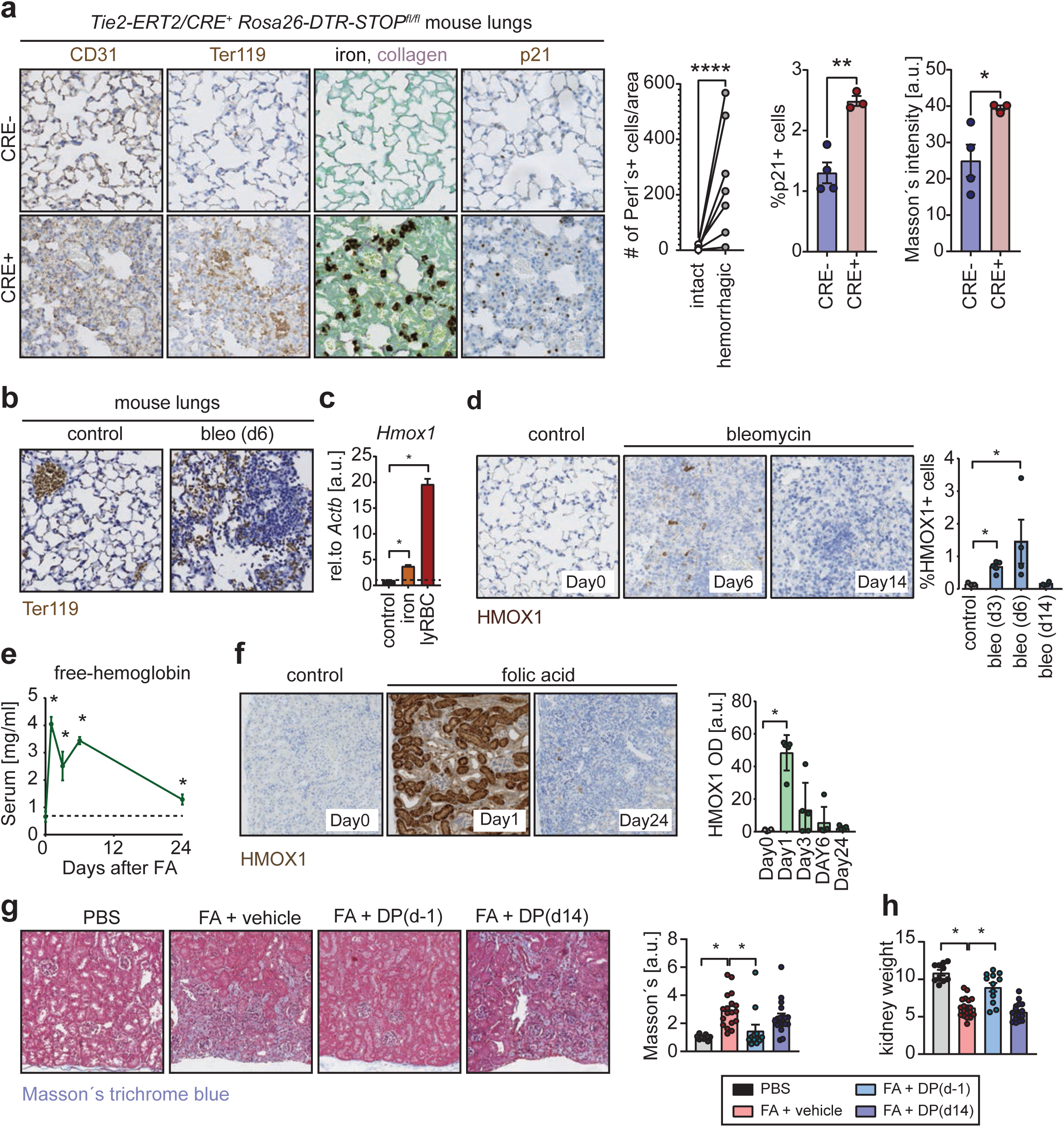
Vascular and hemolytic insults provoke iron accumulation, senescence and fibrogenesis. **a**, Histochemistry for iron and collagen, by combining enhanced Perl’s prussian blue and Sirius Red/Fast Green staining, and immunohistochemistry for the endothelial cell marker CD31, the RBC marker Ter-119, and the senescence marker p21. Representative images of mouse lung sections from *Tie2-ERT2/CRE^-^*(n=3) and *Tie2-ERT2/CRE^+^* (n=3) *Rosa26-DTR- STOP^fl/fl^*mice after two weeks of treatment with intraperitoneally administered tamoxifen and diphtheria toxin (left panel). Number of enhanced Perĺs positive cells per unit area in foci without and with microhemorrhages (middle panel). Percentage of p21^+^ cells and Masson’s Trichrome blue intensity in *Tie2-ERT2/CRE^-^* (n=3) and *Tie2-ERT2/CRE^+^* (n=3) *Rosa26-DTR-STOP^fl/fl^*mice. **b**, Immunohistochemistry for the RBC marker Ter-119. Representative images of mouse lung sections 6 days post-intratracheal administration of PBS (control) or bleomycin. **c**, mRNA levels of *Hmox1* relative to *Actb* in MEFs after 3 days of culture in control conditions, in the presence of iron (660 μM), or in the presence of lysed RBCs (25-fold dilution). **d**, Immunohistochemistry for HMOX1. Representative images of mouse lung sections 0 days (control; n=3), 6 days (n=4), 14 days (n=4) post-intratracheal administration of bleomycin (left panel), and quantification of percentage of HMOX1^+^ cells (right panel). **e**, Free-hemoglobin levels, measured by ELISA, in the sera of mice 0 days (n=7), 1 day (n=5), 3 days (n=5), 6 days (n=5) and 24 days (n=5) after intraperitoneal folic acid. **f**, Immunohistochemistry for HMOX1. Representative images of mouse kidney sections 0 days (n=4), 1 day (n=5), 3 days, 6 days and 24 days (n=5) post-intraperitoneal administration of folic acid (left panel), and quantification of optical density (OD) of HMOX1 staining (right panel). **g-h**, Kidneys from control mice (PBS injected; n=10), and mice in which kidney fibrosis was induced by intraperitoneal folic acid. Mice in the folic acid group either vehicle (n=18) or deferiprone injected daily starting 1 day before (DP(d-1); n=16) or 14 days after (DP(d14); n=18) disease induction by folic acid. Mice were analyzed on 28 post-FA. **g**, Representative images (left panel), and quantification of Masson’s trichrome blue staining. **g**, Kidney weight relative to tibia length. Bar graphs represent mean ± s.e.m.; individual values for each mouse are represented as points; we used two-tailed unpaired t-test. *p < 0.05; **p < 0.01; ***p < 0.001; ****p < 0.0001.

We wondered if vascular or hemolytic injuries are involved in the pathobiology of folic acid induced kidney injury and bleomycin induced lung injury. In the case of bleomycin, we found evidence for microhemorrhagic injuries in the lungs post-bleomycin, as indicated by foci with accumulating extravascular Ter-119^+^ RBCs (**Fig. 3b**). We tested the effects of these microhemorrhagic injuries, by analyzing the levels of heme oxygenase 1 (HMOX1), a gene which we found to be highly responsive to damaged RBCs *in vitro* (**Fig. 3c**). Numbers of HMOX1^+^ cells peaked on day 6 post-bleomycin, and levels normalized by day 14 post-injury (**Fig. 3d**). In the case of folic acid (FA), we found that this compound is hemolytic when applied to isolated RBCs in the concentration range used to provoke kidney injury (**Extended Data Fig. 3b**). *In vivo*, FA- induced hemolysis was reflected by a rapid spike in serum levels of free-hemoglobin (**Fig. 3e**), and in hepatic mRNA levels of haptoglobin (*Hp*, hemoglobin scavenger) and hemopexin (*Hpx*, heme scavenger) (**Extended Data Fig. 3c and 3d**), reaching their maximum as early as one day post-injury, which was followed by a consecutive decline in these effects. In line with these findings, FA induced a transient and robust increase in the levels of heme oxygenase 1 (HMOX1) in the kidneys one day post-injury (**Fig. 3f and Extended Data Fig. 3e**), possibly in response to the hemolytic filtrate. Of note, all the parameters measuring RBC damage were consistent with a rapid but transient release of hemolytic iron, which is in line with the kinetics of hepatic hepcidin (**see above Fig. 1i,j**).

Finally, to test the causal role of iron in FA-induced renal fibrosis, we treated mice with folic acid to provoke kidney injury in the presence or absence of the iron chelator deferiprone. In one group of mice, labeled DP(d-1), deferiprone was administered starting one day before FA and continued until analysis at day 28 post-FA. In another group of mice, DP(d14), treatment was initiated at day 14 post-FA until day 28. Interestingly, mice in the DP(d-1) group were largely protected from FA induced kidney fibrosis and atrophy (**Fig. 3g,h**), however, deferiprone was not able to reverse already established fibrotic scars (**Fig. 3g,h and Extended Data Fig. 3f**). We conclude that vascular injuries and hemolytic RBCs have the capacity to drive iron accumulation, senescence, and fibrosis.

### Iron and damaged RBCs cause cellular senescence *in vitro*

To gain mechanistic insight into the connection between iron and senescence, we explored the effects of iron on *in vitro* cultured cells. We found in pilot experiments that for iron uptake to occur, cells had to be primed first by culturing them in low transferrin-bound iron conditions (achieved by reducing the serum to 0.5%). Under these conditions, addition of iron, lysed RBCs, or hemin (a derivative of the iron-bound cofactor of hemoglobin), efficiently triggered senescence (measured by senescence-associated beta-galactosidase, SA-β-GAL) in all the cell lines tested, namely, primary mouse embryo fibroblasts (MEFs), immortalized mouse endothelial cells (H5V), non- immortalized human lung fibroblasts (IMR90), and non-immortalized human umbilical vein endothelial cells (HUVEC) (**Fig. 4a**). This response was studied in further detail in MEFs. First, we observed that iron and lysed RBCs upregulated the transcription of ferritin subunits (*Fth1* and *Ftl*) (**Fig. 4b**), suggesting increased intracellular iron storage, and prominent SASP cytokines *Il6* and *Ccl2* (**Fig. 4c and 4d**). In the case of lysed RBCs, these transcriptional effects were ablated by the presence of deferiprone (**Fig. 4d**), indicating that iron is the key mediator. Iron also caused a dramatic elevation in cellular ROS levels (**Fig. 4e**) and cells presented with an expansion in their lysosomal mass, as measured by lysotracker staining (**Fig. 4f**). Interestingly, ROS scavengers tocopherol and ferristatin potently mitigated lysosomal expansion in response to iron (**Fig. 4f**). Similar observations were made in human melanoma cells SK-MEL-103 (**Fig. 4g**). ROS scavengers also prevented the emergence of large SA-β-GAL positive cells (**Fig. 4h**) and so was hypoxia (**Extended Data Fig. 4a**). We conclude that iron, in its free form or when released from damaged RBCs, is a potent trigger of cellular senescence *in vitro*.

**Fig. 4.**
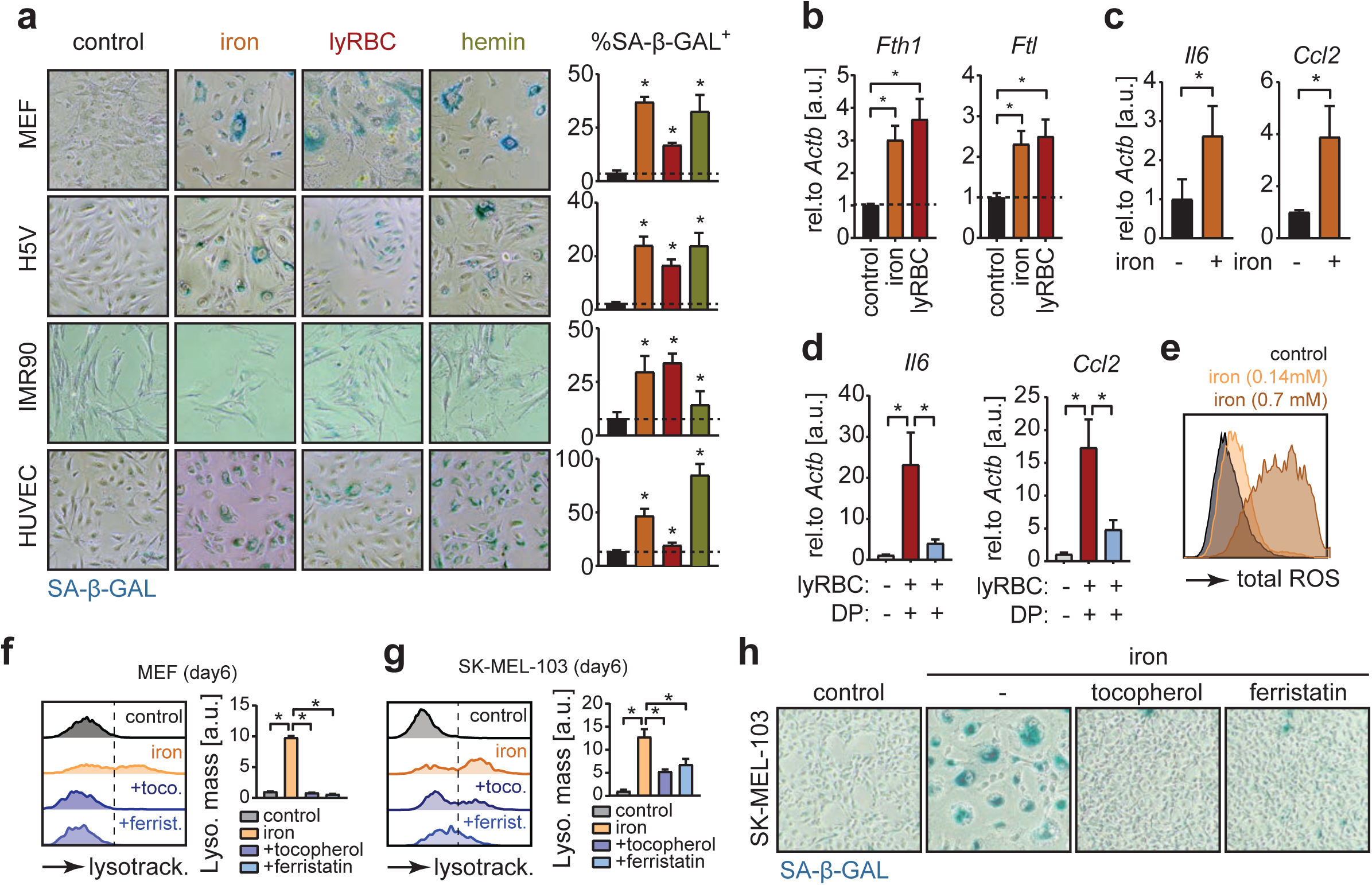
Iron and damaged RBCs cause cellular senescence *in vitro*. **a**, SA-β-GAL staining of H5V mouse endothelial cells, IMR90 human lung epithelial fibroblasts, HUVEC, human umbilical vein endothelial cells, and MEFs, which were cultured for one day in RPMI medium containing 0,5% FBS (to prime cells for iron uptake), followed by 3 days of culture in the same medium supplemented with vehicle (control), iron (660 μM), lysed RBCs (25-fold dilution), or hemin (10 μM), and consecutive culture for 2 weeks in complete medium. Representative images (left panel). Percentage of SA-β-GAL^+^ cells (right panel). **b**, mRNA levels of *Fth1* and *Ftl* relative to *Actb* in MEFs, which were cultured for one day in RPMI medium containing 0,5% FBS (to prime cells for iron uptake), followed by 3 days of culture in the same medium supplemented with vehicle (control), iron (660 μM), or lysed RBCs (25-fold dilution). **c**, mRNA levels of *Il6* and *Ccl2* relative to *Actb* in MEFs, which were cultured for one day in RPMI medium containing 0,5% FBS (to prime cells for iron uptake), followed by 3 days of culture in the same medium supplemented with vehicle (control), or iron (660 μM). **d**, mRNA levels of *Il6* and *Ccl2* relative to *Actb* in MEFs which were cultured for one day in RPMI media containing 0,5% FBS (to prime cells for iron uptake), followed by 3 days of culture in the same medium supplemented with vehicle (control), or lysed RBCs (25- fold dilution) with or without deferiprone (1 mM). **e**, Flow-cytometric analysis of total-ROS levels in MEFs after 7 days in control conditions or in the presence of iron (140 μM or 660 μM). **f-g**, Flow-cytometric analysis of lysosomal mass by lyso-tracker staining of **f**, MEFs and **g**, SK-MEL- 103 cells after culturing cells for 6 days in control conditions or in the presence of the ROS scavengers tocopherol (50 μM) and ferristatin (1 μM). Representative histograms (left panel). Quantification of lysosomal mass by normalizing the delta mean fluorescence intensity (ΔMFI) to control values. **a**, SA-β-GAL staining of SK-MEL-103 cells after two weeks of culture in control conditions, or in the presence of iron with or without tocopherol (50 μM) or ferristatin (1 μM). Bar graphs represent mean ± s.e.m.; we used two-tailed unpaired t-test. *p < 0.05; **p < 0.01; ***p < 0.001; ****p < 0.0001.

### Cellular damage triggers iron accumulation that fuels ROS and the SASP

We were intrigued by the different kinetics of injury-induced iron release and injury-induced iron accumulation. Specifically, tissue injuries provoked a transient burst of hemolysis (detectable by HMOX1 and by hepatic *Hamp*, *Hp*, and *Hpx*), while iron accumulation in the fibrotic lesions was progressive and continuous even after hemolysis subdued to basal levels. We wondered if progressive iron accumulation could be an intrinsic property of damaged cells, even in the absence of an initial burst of extracellular iron. For this, we investigated the kinetics of iron accumulation after cellular damage *in vitro* (bleomycin, doxorubicin, irradiation and palbociclib) under normal culture conditions. We observed a progressive increase in total iron levels and upregulation of ferritins as early as 3 days post-insult (**Fig. 5a,b and Extended Data Fig. 5a**), suggesting that iron accumulation starts before the establishment of senescence. Using enhanced Perl’s prussian blue staining, we found that iron deposits concentrated in cytoplasmic puncta in senescent cells (**Fig. 5c,d and Extended Data Fig. 5b,c**). Transmission electron microscopy (TEM) suggested that these puncta correspond to lysosomes containing abundant ferritin-bound iron (**Fig. 5e,f**). In addition to increased ferritin-bound iron (inert Fe^3+^), we also found that the labile iron (oxidative Fe^2+^) pool was significantly elevated in senescent cells (**Fig. 5g,h**).

**Fig. 5.**
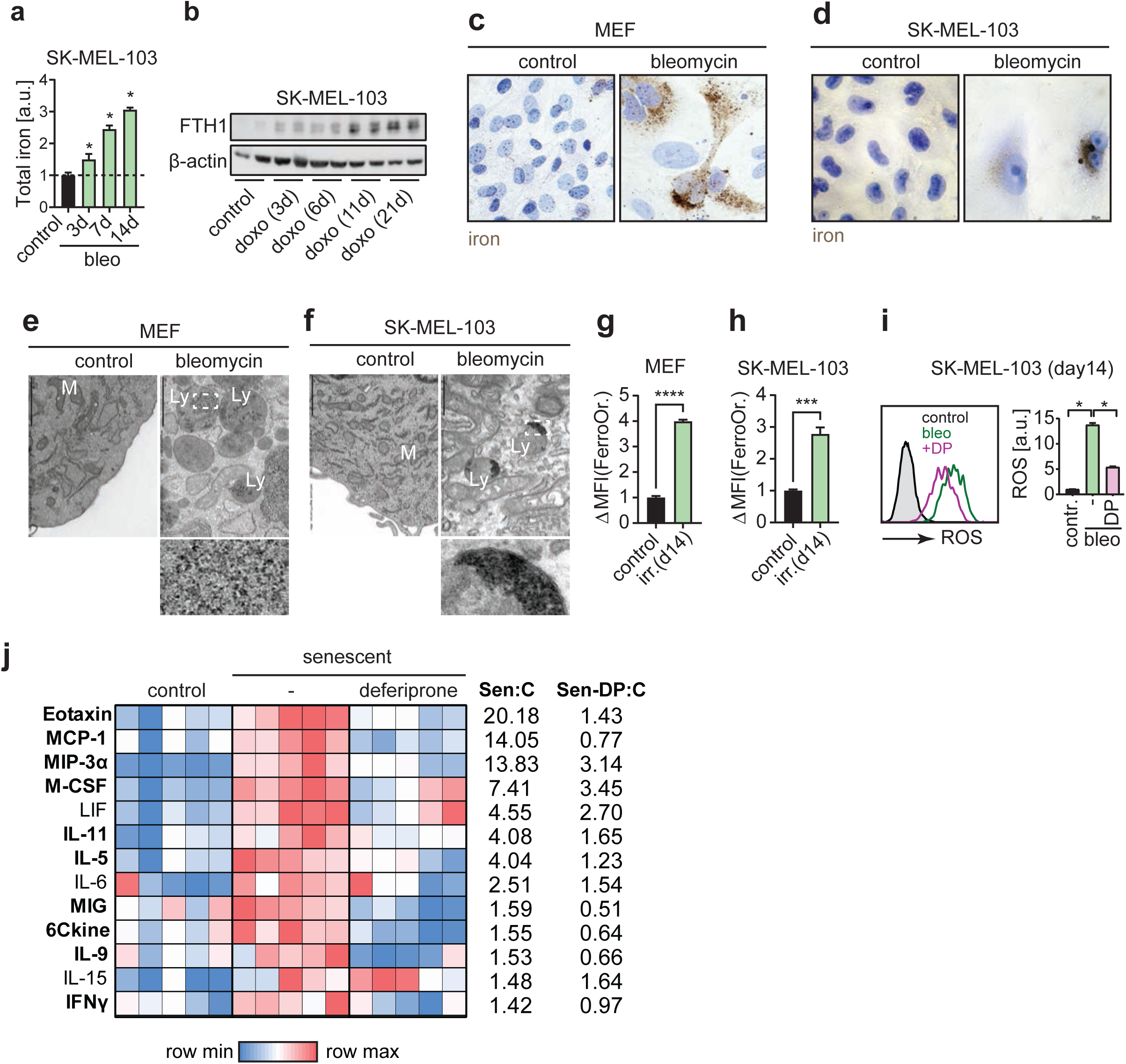
Cellular damage triggers iron accumulation that fuels ROS and the SASP. **a**, Total iron levels in SK-MEL-103 cells before and 3 days, 7 days, and 14 days post-damage by bleomycin. **b**, Ferritin heavy chain (FTH1) levels in SK-MEL-103 cells before and 3 days, 6 days, 11 days and 21 days post-damage by doxorubicin. **c-d**, Enhanced Perl’s prussian blue staining of **c**, MEFs and **d**, SK-MEL-103 cells, controls and 21 days post-damage by bleomycin. **e-f**, Transmission electron microscopical images of **e**, MEFs and **f**, SK-MEL-103 cells, controls and 21 days post-damage by bleomycin. Inserts show accumulation of electron-dense material in the lysosomes compatible in size and organization with ferritin-bound iron. Lysosomes (Ly); Mitochondria (M). **g-h**, Levels of labile iron (Fe^2+^) in control and senescent (irradiation-induced) **g**, MEFs and **h**, SK-MEL-103 cells, measured by flow-cytometry using the (Fe^2+^)-specific dye FerroOrange. **i**, Levels of total-ROS in control and senescent (bleomycin-induced) SK-MEL-103 cells, which received 30 minutes prior to the measurement vehicle or deferiprone (DP; 200 μM). Total-ROS levels measured by flow-cytometry. Representative histograms (left panel), and quantification (right panel). **j**, Heatmap showing protein levels of cytokines and chemokines in control, and senescent MEFs, and in senescent MEF cultured for three days in the presence of Deferiprone (DP; 100 μM). Cytokines listed were all significantly induced in senescent MEFs, when compared to controls. Cytokines marked by bold fonts were significantly suppressed by deferiprone. Fold change relative to the controls for senescent MEFs is listed in the column “Sen:C”, for senescent MEFs treated with deferiprone is listed in the column “Sen-DP:C”. In all cases, bar graphs represent mean ± s.e.m.; we used two-tailed unpaired t-test. *p < 0.05; **p < 0.01; ***p < 0.001; ****p < 0.0001.

We wondered if the high ROS levels characteristic of senescent cells^47, 48^ are fueled, at least in part, by labile iron. Interestingly, when fully senescent cells were treated for 30 min with deferiprone (DP), ROS levels were significantly reduced (**Fig. 5i and Extended Data Fig. 5d,f**). ROS is known to be a main trigger of the secretory phenotype of senescent cells (SASP)^49, 50^ and, therefore, we asked if iron chelation would also interfere with the SASP. Control and fully senescent cells were tested for the intracellular protein levels of a total of 44 cytokines and chemokines, out of which 13 were significantly elevated upon senescence (**Fig. 5j**). Interestingly, treating senescent cells with deferiprone for three days ablated the large majority of cytokines and chemokines in the SASP (10 out of 13 were significantly suppressed) (**Fig. 5j**). We conclude that cellular damage triggers rapid and progressive iron accumulation even in the absence of an excess of extracellular iron. Moreover, the resulting intracellular accumulation of iron fuels the generation of high levels of ROS and the SASP.

### Non-invasive assessment of fibrosis by MRI-based detection of iron

We wondered whether iron accumulation in fibrotic tissues could be used as a proxy to detect fibrosis using non-invasive medical imaging. Prior to this, we documented that the percentage of Perĺs positive cells was directly correlated with the Massońs trichrome blue staining (**Fig. 6a**) and inversely correlated with kidney weight (**Fig. 6b**) in mice treated with FA. Of note, the majority of Perĺs positive cells were located cortically in the FA-treated kidneys, while the medullae were largely negative (**Extended Data Fig. 6a**). Iron can be detected by magnetic resonance imaging (MRI) using the R2* relaxation mapping^51^. Similar to the histological observations by Perĺs staining, R2* mapping detected a significantly elevated signal in the cortex, but not in the medulla, of the kidneys of FA-treated mice (**Fig. 6c,d**). The R2* signals in the cortices of individual kidneys correlated positively with their corresponding Perĺs (**Fig. 6e**) and Massońs staining (**Fig. 6f**), and correlated negatively with kidney weight (**Fig. 6g**).

**Fig. 6.**
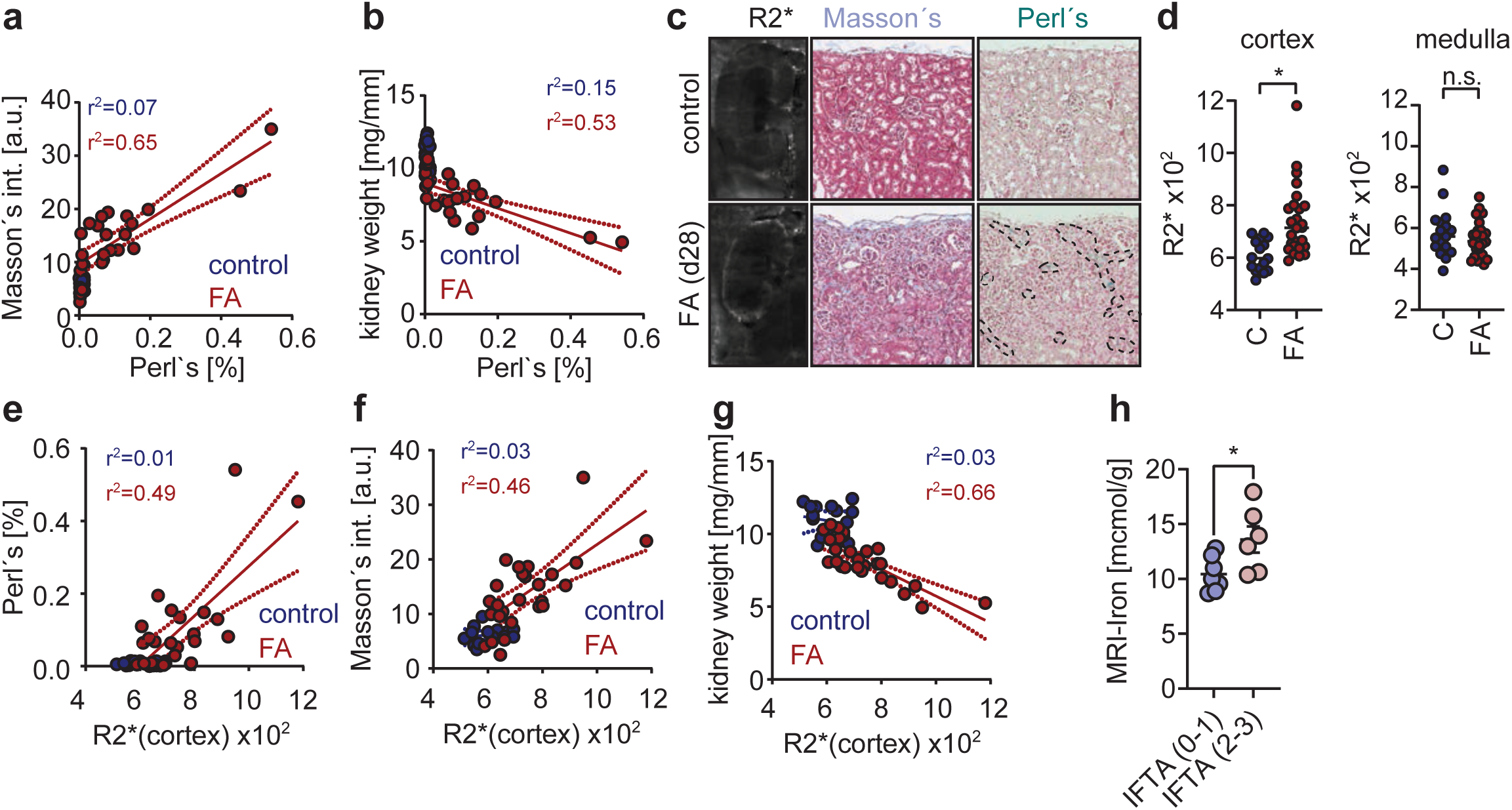
Non-invasive assessment of fibrosis by MRI-based detection of iron. **a-b**, Correlation analysis between **a**, Perl’s and Masson’s trichrome blue staining intensity, and **b**, Perl’s staining intensity and kidney weight in mice. Each dot represents a mouse kidney from mice 28 days post- intraperitoneal PBS (blue; n=16) or intraperitoneal folic acid (red; n=26). **c**, Representative images of R2* map acquired by MRI, Masson’s trichrome staining, and Perĺs staining of kidneys from mice 28 days after injection with PBS or folic acid. **d**, Mean R2* value in the kidney cortex and in the kidney medulla of mice 28 days after treatment with intraperitoneal injection of PBS or folic acid. **e-g**, Correlation analysis between the mean R2* value measured in the kidney cortex and **e,** intensity of Perl’s staining in the kidney cortex, **f,** intensity of Masson’s staining in the kidney cortex, and **g**, kidney weight. Each dot represents a mouse kidney from mice receiving intraperitoneal PBS (blue; n=16) or intraperitoneal folic acid (red; n=26) analyzed 28 days after injection. **h**, R2* values in the kidney cortex were measured in kidney allograft recipient patients (n=3). We compared R2* values obtained in patients with no or moderate fibrosis (IFTA (0-1); n=7) to patients with pronounced fibrosis (IFTA (2-3); n=6). Bar graphs represent mean ± s.e.m.; we used two-tailed unpaired t-test. *p < 0.05; **p < 0.01; ***p < 0.001; ****p < 0.0001.

To evaluate if this method can be utilized in a clinical setting, we performed a pilot study on patients with diabetic kidney disease. We analyzed 13 patients by MRI for their R2* signal in the kidney cortex, and for their level of fibrosis measured on patient biopsies (IFTA score, interstitial fibrosis and tubular atrophy). We found that patients with high IFTA score (2 and 3) presented with significantly higher R2*, than patients with low IFTA score (0 and 1) (**Fig. 6h**). Collectively, we conclude that iron measurement by MRI may be a clinically translatable tool for non-invasive detection of fibrotic tissue remodeling.

## DISCUSSION

Iron accumulation in the context of fibrosis has been reported previously^30,31,32, 52^, here we showed for the first time that this is not dependent on a specific organ or insult, but is an early event upon tissue injury that, once initiated, progresses and becomes a defining feature of fibrotic diseases. We demonstrate that exposure to iron-overload, such as caused by direct administration of iron, or vascular or hemolytic insults, is sufficient to initiate fibrogenesis and senescence. However, we also show that the accumulation of iron in fibrotic lesions continues even when the initial burst of extracellular iron caused by tissue injury has subdued. This has led us to discover that iron accumulation is an intrinsic feature of damaged cells regardless of the type of damaging insult that does not require extracellular iron excess. Therefore, for the purposes of this discussion, it is helpful to distinguish between (1) the role of “excessive extracellular iron” as a potential pathophysiological trigger of senescence and fibrosis and (2) the role of “excessive intracellular iron” as a driver of the pathological effects of senescent cells through the SASP.

Regarding the role of “excessive extracellular iron”, there is abundant evidence linking hereditary and acquired hemochromatosis to fibrotic diseases^25,26,27,29^. Our data extend this concept to fibrotic diseases unrelated to extracellular iron overload. Indeed, many fibrotic processes are of unknown etiology. We present data indicating that small vascular injuries with associated iron release could underlie some fibrotic diseases. Indeed, in the case of systemic sclerosis, a disease characterized by multi-organ fibrosis, vascular insufficiency manifests early in the disease, and its potential involvement in the disease pathobiology has been proposed^53, 54^. Patients with complement mediated hemolytic anemia are also at risk of developing fibrosis in their kidneys, a condition known as hemolytic uremic syndrome^55^. In addition, we found that iron potently suppresses VEGF levels, and drives capillary rarefaction. VEGF is known for its effects to promote blood flow in the local circulation^56^, and it is attractive to speculate that perhaps vascular rarefaction as observed in fibrotic conditions^11,12,13^, is an attempt by the body to limit bleeding through the suppressive effect of vascular origin iron on VEGF.

Regarding “excessive intracellular iron”, we report that this is initiated by cellular damage and progressively increases together with the development of the senescent phenotype. This process does not require the presence of abnormal amounts of extracellular iron. Senescent cells upregulate the expression of ferritin subunits and accumulate large amounts of ferritin-bound iron in lysosomes. In addition, senescent cells also have abnormally high levels of labile iron, which can be a main cause of ROS. Indeed, treatment of senescent cells with an iron chelator strongly reduces their ROS levels. Previous reports have connected ROS with the secretory phenotype of senescent cells^49, 50^. Based on this, we also show that iron chelation efficiently suppresses the SASP, thereby mechanistically connecting damage-initiated iron uptake with ROS and the SASP. The intrinsic and cell-autonomous accumulation of intracellular iron by damaged and senescent cells explains why, during fibrogenesis, iron continues accumulating even in the presence of normal extracellular iron levels.

Finally, we demonstrate in a proof of principle study that iron detection by MRI could be a useful diagnostic tool to identify fibrotic remodeling early. MRI based iron detection has been first used in a clinical setting in a seminal study on thalassemia major patients to detect iron accumulation in the heart^51^. Later studies demonstrated that MRI based iron detection allows for assessment of iron deposition in the liver, spleen and other organs, and thereby for the clinical assessment of patients with hereditary hemochromatosis or hemoglobinopathies^57, 58^. Here, for the first time, we show that the same technology can be applied to assess fibrotic burden non- invasively in mice and patients with kidney fibrosis. These individuals have no genetic or acquired iron accumulation syndrome, but accumulate iron in their kidneys in association with fibrosis. We find that iron levels correlate with the grade of the fibrosis and with kidney atrophy, and urge other clinicians to test in the context of fibrotic diseases the same method for its utility to improve early detection of fibrotic disease.

## ACKNOWLEDGEMENTS

We thank the TEM-SEM Electron microscopy Unit from Scientific and Technological Centers (CCiTUB), Universitat de Barcelona, and their staff for their support and advice on TEM technique. We are thankful to the Magnetic Resonance Imaging Core Facility of the Institut d’Investigacions Biomèdiques August Pi i Sunyer (IDIBAPS) for the scientific and technical support in MRI acquisition and analysis. Work in the laboratory of M.S. was funded by the IRB and “laCaixa” Foundation, and by grants from the Spanish Ministry of Science co-funded by the European Regional Development Fund (ERDF) (SAF2017-82613-R), European Research Council (ERC- 2014-AdG/669622), and Secretaria d’Universitats i Recerca del Departament d’Empresa i Coneixement of Catalonia (Grup de Recerca consolidat 2017 SGR 282). M.M. received funding from the European Union’s Horizon 2020 research and innovation programme under the Marie Sklodowska-Curie grant agreement (No 794744) and from the Spanish Ministry of Science (RYC2020-030652-I /AEI /10.13039/501100011033). Further funding includes (FPU- 18/05917) to V.P.L. PID2021-122436OB-I00 from MCIN/ AEI /10.13039/501100011033 / FEDER, UE, and grant RETOS COLABORACION RTC2019-007074-1 from MCIN/AEI /10.13039/501100011033 to M.S. Instituto de Salud Carlos III through the projects PI18/00910 and PI21/00931 (Co-funded by European Regional Development Fund. ERDF, a way to build Europe). We thank CERCA Programme / Generalitat de Catalunya for institutional support. To J.M.C. Individual Fellowship of the German Cardiac Society, Individual Fellowship of the German Research Foundation and Juan de la Cierva Fellowship to K.M. Instituto de Salud Carlos III (PI 20/01360, FEDER funds) to G.A.

We are thankful to Dr. Daniel Muñoz Espin for sending us the IMR90 cells stably transduced with a tamoxifen inducible Ras-G12V.

## EXTENDED DATA

**Extended Data Fig. 1.**
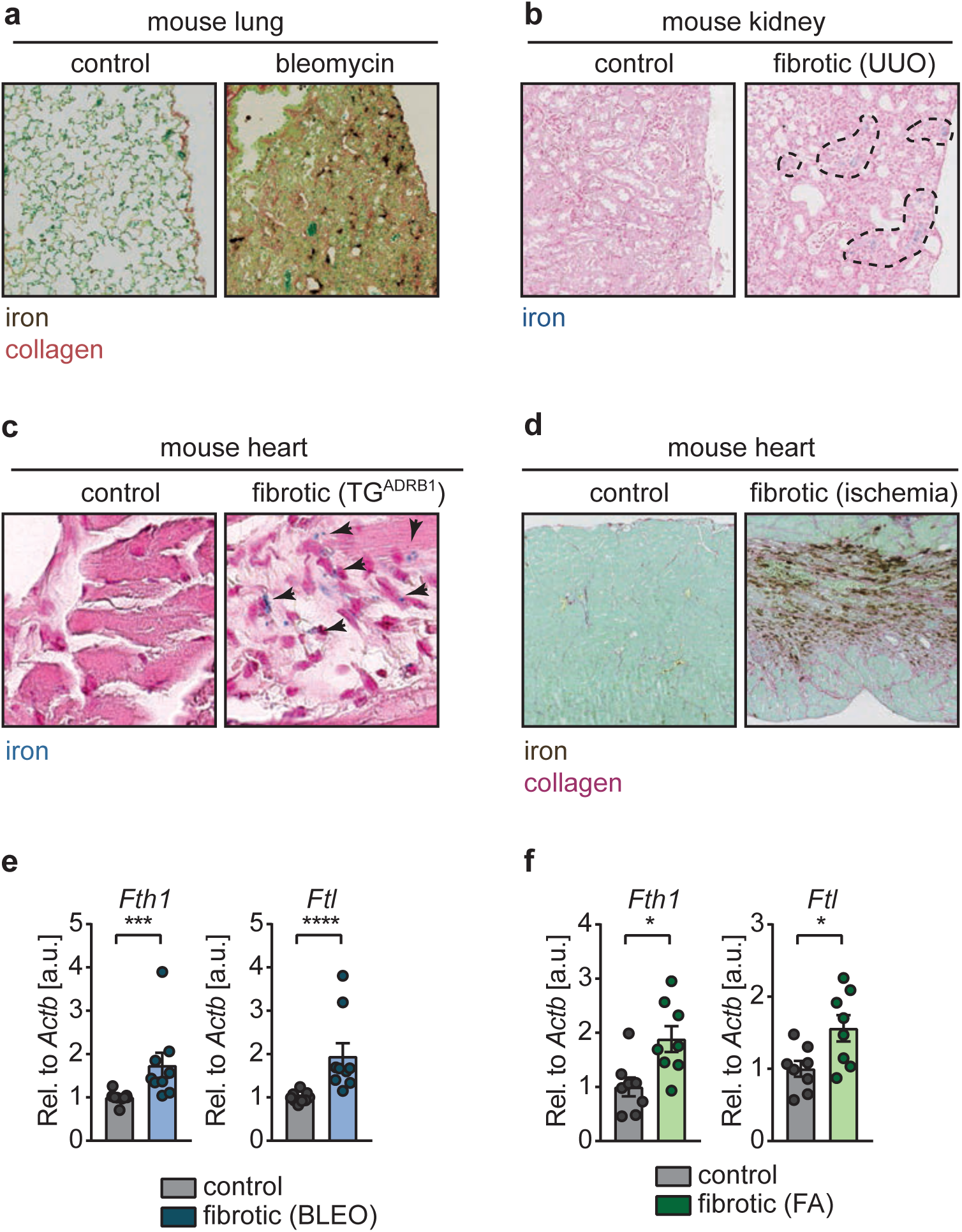
Tissue injury provokes iron accumulation that progresses throughout fibrosis. **a**, Histochemistry for iron and collagen by combining enhanced Perl’s prussian blue and Sirius Red/Fast Green staining. Representative images of mouse lung sections 14 days post-intratracheal administration of PBS or bleomycin. **b**, Histochemistry for iron by regular Perl’s prussian blue staining. Representative images of kidneys from control mouse and mouse in which kidney fibrosis was provoked by unilateral ureteral obstruction. **c**, Histochemistry for iron by regular Perl’s prussian blue staining. Representative images of mouse hearts from wild-type mice (control) and mice with transgenic overexpression of adrenergic receptor beta (fibrotic, *TG^ADRB^*^1^). **d**, Histochemistry for iron and collagen by combining enhanced Perl’s prussian blue and Sirius Red/Fast Green staining. Representative images of mouse heart sections undergoing sham surgery (control), and from a mouse in which fibrosis was provoked by ischemic coronary artery ligation. **e-f**, mRNA levels of *Fth1* and *Ftl* relative to *Actb* **e**, in mouse lungs 14 days post-intratracheal administration of PBS (control; n=8) or bleomycin (fibrotic, BLEO; n=8), and **f**, in mouse kidneys 28 days post-intraperitoneal administration of PBS (control; n=8) or folic acid (fibrotic, FA; n=8). Bar graphs represent mean ± s.e.m.; individual values for each mouse are represented as points; we used two-tailed unpaired t-test. *p < 0.05; **p < 0.01; ***p < 0.001; ****p < 0.0001.

**Extended Data Fig. 2.**
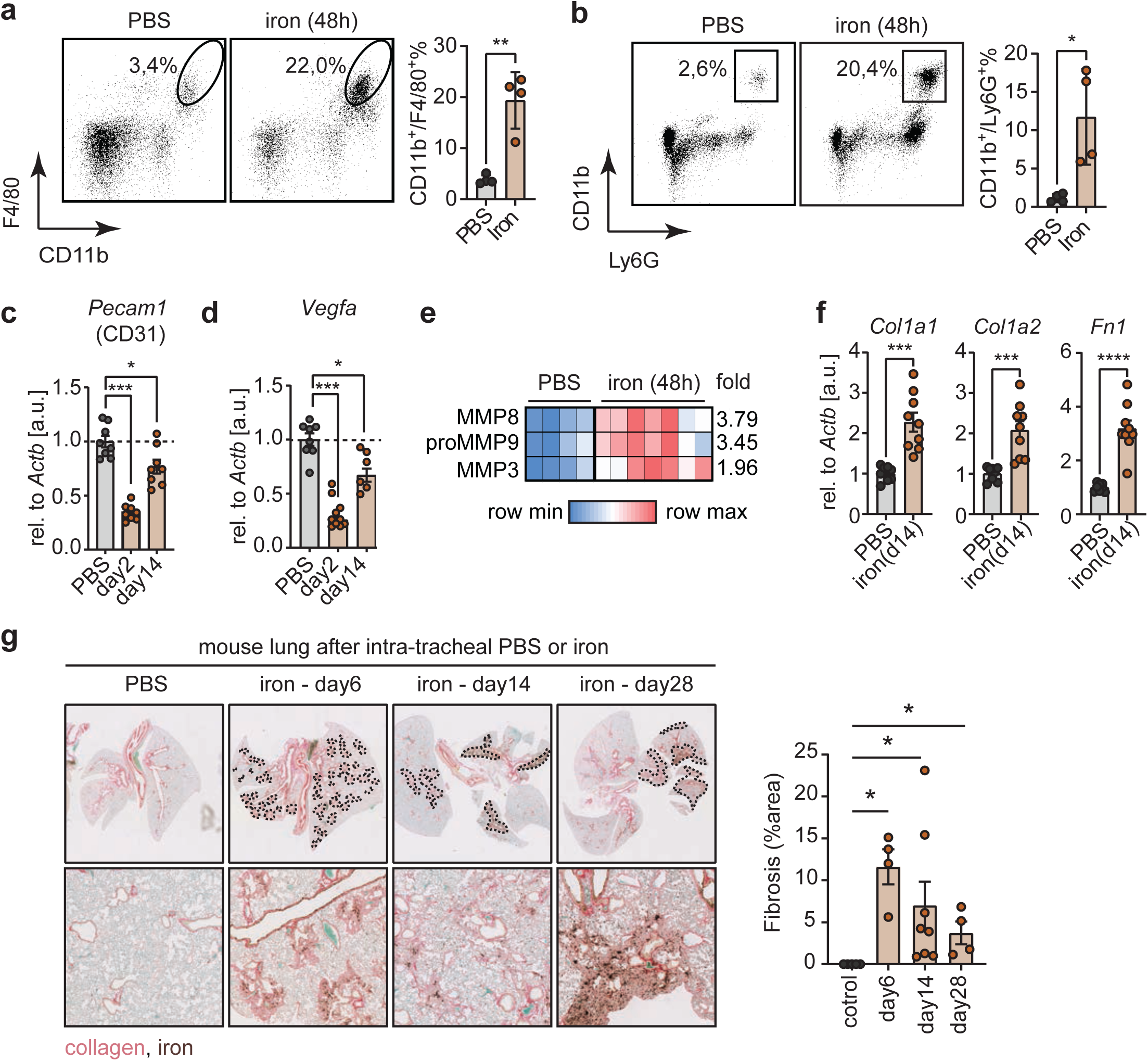
Iron initiates fibrogenesis. **a-b**, Flow-cytometric analysis of dissociated lung tissues for the **a**, macrophage markers CD11b and F4/80, and **b**, the neutrophil markers CD11b and Ly6G. Representative dot plots of lung cells from mice 2 days post-intratracheal delivery of PBS (n=4) or iron (500 nM; n=4), and quantification. **c-d**, mRNA levels of **c**, *Pecam1* (CD31) and **d**, *Vegfa* relative to *Actb* in mouse lungs 2 days post-intratracheal administration of PBS (n=8) or 2 days (n=8-10) and 14 days (n=6-8) post-intratracheal administration of iron (500 nM). **d**, Heatmap of significantly altered matrix metalloproteinases (MMP) in the lungs of mice 2 days post-intratracheal delivery of PBS (n=4) or iron (500 nM; n=7). Lung lysates were tested by a multiplex protein assay for 5 MMPs out of which 3 were significantly changed, with a fold-change as shown to the right. **f**, mRNA levels of *Col1a1*, *Col1a2*, *and Fn1* relative to *Actb* in mouse lungs 14 days post-intratracheal administration of PBS (n=8) or iron (500 nM; n=9). **f**,Histochemistry for iron and collagen, by combining enhanced Perl’s prussian blue and Sirius Red/Fast Green staining. Representative images of mouse lung sections from mice 2 days post-intratracheal administration of PBS (control; n=5) or 6 days (n=4), 14 days (n=8), and 28 days (n=4) after intratracheal administration of iron (500 nM; n=4), and quantification. Dashed lines delineate iron and collagen rich areas. Bar graphs represent mean ± s.e.m.; individual values for each mouse are represented as points; we used two-tailed unpaired t-test. *p < 0.05; **p < 0.01; ***p < 0.001; ****p < 0.0001.

**Extended Data Fig. 3.**
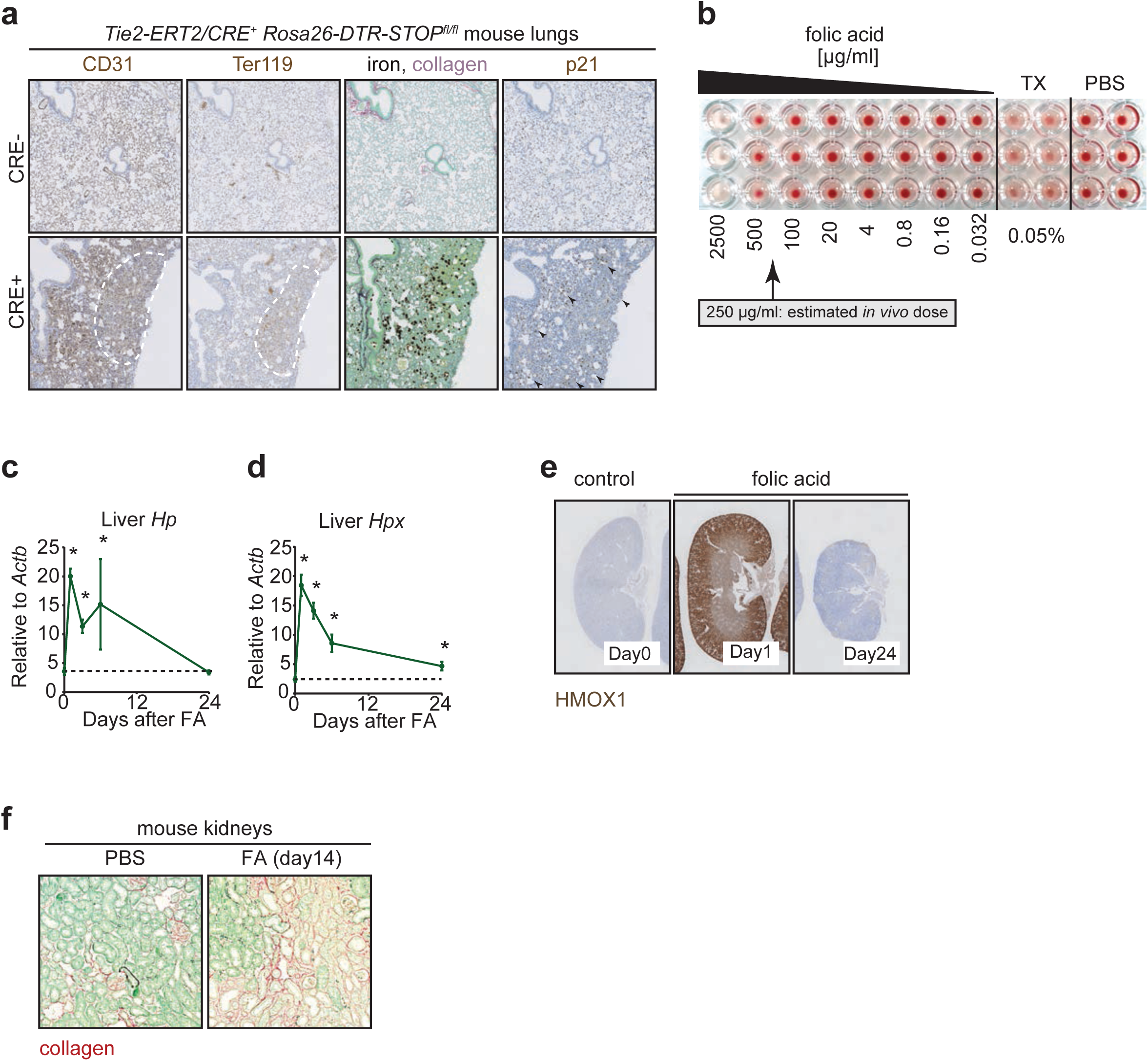
Vascular and hemolytic insults provoke iron accumulation, senescence and fibrogenesis. **a**, Histochemistry for iron and collagen, by combining enhanced Perl’s prussian blue and Sirius Red/Fast Green staining, and immunohistochemistry for the endothelial cell marker CD31, the RBC marker Ter-119, and the senescence marker p21. Representative images of mouse lung sections from *Tie2-ERT2/Cre^-^* and *Tie2-ERT2/CRE^+^ Rosa26-DTR-STOP^fl/fl^* mice after two weeks of treatment with intraperitoneally administered tamoxifen and diphtheria toxin. Dashed line indicates area with reduced CD31 expression associated with accumulation of micro-hemorrhagic RBCs. **b**, Ex vivo RBC hemolysis assay with different doses of folic acid, with triton X-100 as a positive control (TX) as a positive control and PBS as a negative control. **c-d**, mRNA levels of **c**, haptoglobin (*Hp*) and **d**, hemopexin (*Hpx*) relative to *Actb* in the livers of mice from before (n=6), and 1 day (n=5), 3 days (n=5), 6 days (n=5), or 24 days (n=5) post-folic acid. **f**, Immunohistochemistry for HMOX1. Representative images of mouse kidney sections 0 days, 1 day, 3 days, and 24 days post-intraperitoneal administration of folic acid. **f**, Histochemistry for collagen by Sirius Red/Fast Green staining. Representative images of mouse kidney sections 14 days post-intraperitoneal administration of PBS or folic acid. Bar graphs represent mean ± s.e.m.; individual values for each mouse are represented as points; we used two-tailed unpaired t-test. *p < 0.05; **p < 0.01; ***p < 0.001; ****p < 0.0001.

**Extended Data Fig. 4.**
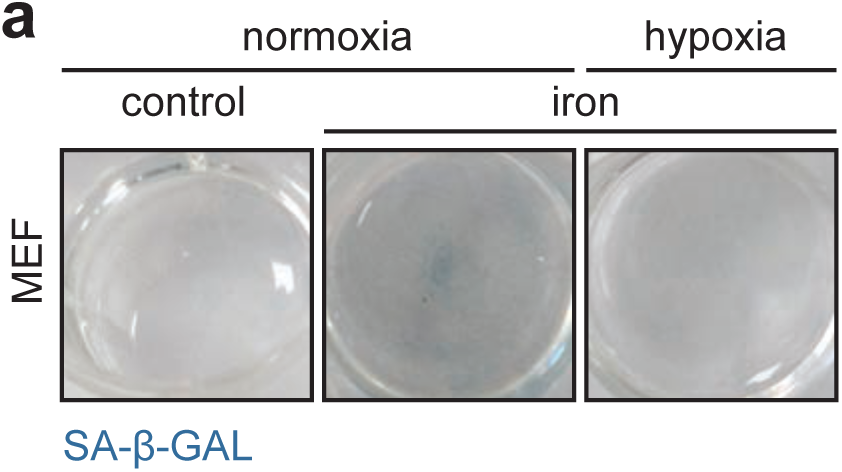
Iron and damaged RBCs cause cellular senescence *in vitro*. **a**, Representative images of culture plate wells confluent with MEFs that were cultured in control conditions or in the presence of iron in normoxia or hypoxia, and subsequently stained for SA-β-GAL.

**Extended Data Fig. 5.**
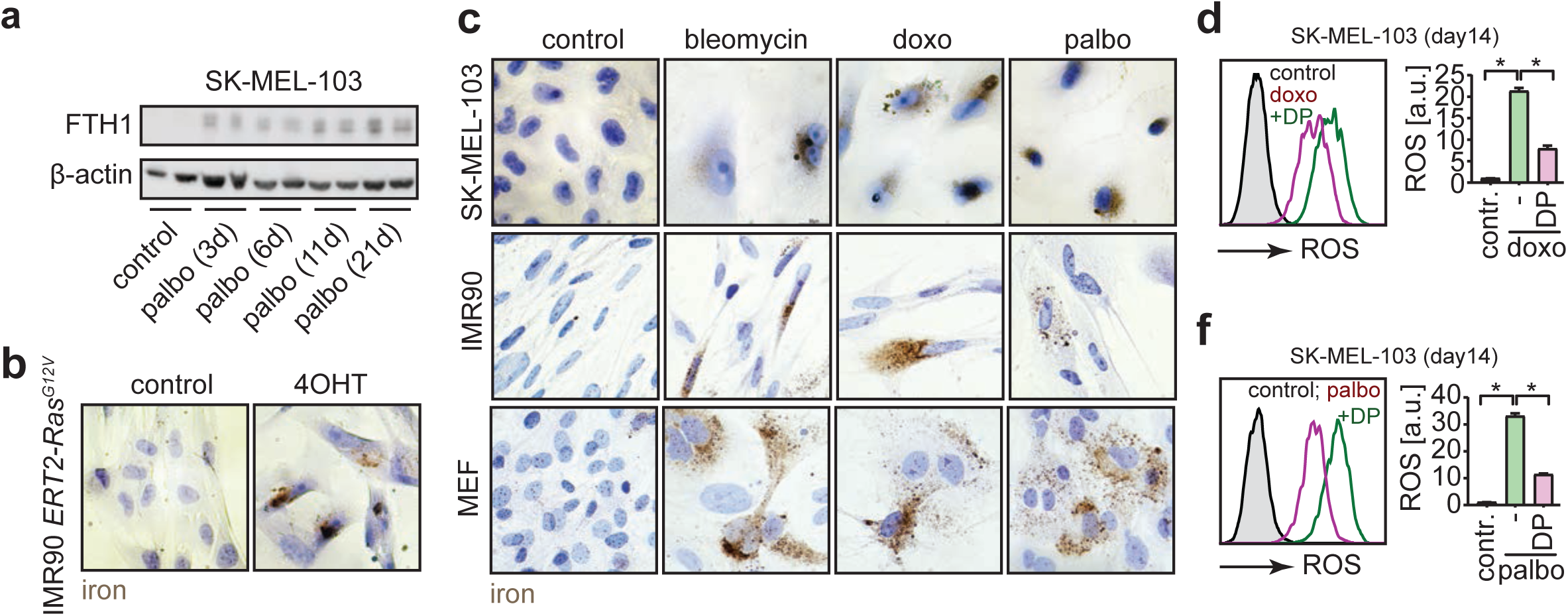
Cellular damage triggers iron accumulation that fuels ROS and the SASP. **a**, Ferritin heavy chain (FTH1) levels in SK-MEL-103 cells before and 3 days, 6 days, 11 days and 21 days post-damage by palbociclib. **b-c**, Enhanced Perl’s prussian blue staining of **b**, IMR90 **c**, SK-MEL-103, IMR90, and MEF control or senescent. Senescence was induced by overexpression of mutant Ras^G12V^, bleomycin, doxorubicin or palbociclib, followed by culture for 21 days. **d-f**, Levels of total-ROS in control and senescent SK-MEL-103 cells, which received 30 minutes prior to the measurement vehicle or deferiprone (DP; 200 μM). Senescence was induced by **d**, doxorubicin, or by **d**, palbociclib. Total-ROS levels measured by flow-cytometry. Representative histograms (left panel), and quantification (right panel). In all cases, bar graphs represent mean ± s.e.m.; we used two-tailed unpaired t-test. *p < 0.05; **p < 0.01; ***p < 0.001; ****p < 0.0001.

**Extended Data Fig. 6.**
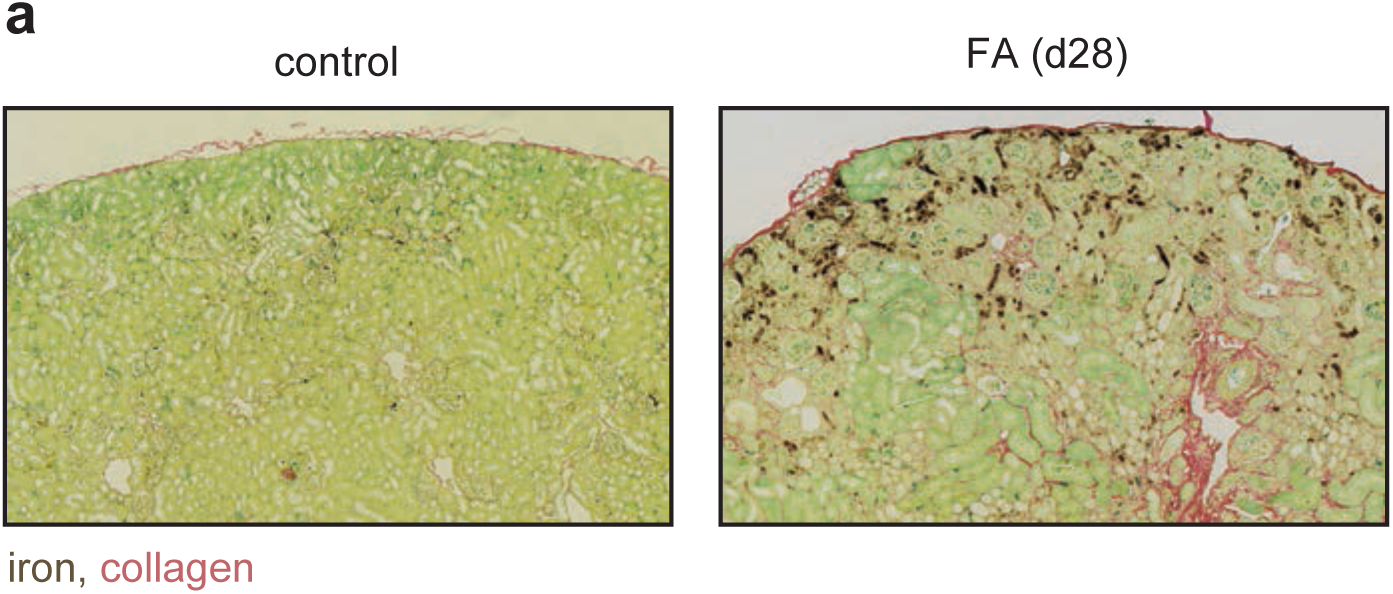
Non-invasive assessment of fibrosis by MRI-based detection of iron. **a**, Histochemistry for iron and collagen by combining enhanced Perl’s prussian blue and Sirius Red/Fast Green staining. Representative images of mouse kidneys

## MATERIALS & METHODS

### Cells

Cell lines SK-MEL-103 (human melanoma), H5V (mouse endothelial cells), HUVEC (human umbilical vein endothelial cells), IMR90 (human fetal lung fibroblast) were obtained from ATCC. Mouse embryonic fibroblasts (MEF) were isolated as described^1^. Cells were cultured in standard DMEM or RPMI supplemented with 10% FBS, unless indicated otherwise, and penicillin–streptomycin (all from Gibco) in 20% O_2_ and 5% CO_2_ at 37°C. Cells were routinely tested for mycoplasma contamination using the mycoplasma tissue culture NI (MTC-NI) Rapid Detection System (Gen-Probe).

### Mice

Mice were housed under specific pathogen free conditions in the mouse facility of the Institute for Research in Biomedicine (IRB) in accordance with the protocols approved by the Animal Care and Use Ethical Committee of animal experimentation of the Barcelona Science Park (CEEA-PCB) and the Catalan Government. The animals were kept under a 12 h-12 h light-dark cycle and allowed unrestricted access to food and water. *TG^Abrd^*^1^ mice, overexpressing the human ß1-adrenoceptor in cardiac myocytes driven by the α-myosin heavy chain promoter, develop spontaneous cardiomyopathy, mimicking human heart failure, were described earlier (Engelhardt et al. 1999). Section of the cardiac tissue from *TG^Abrd^*^1^ mice and their wild-type littermates were kindly sent by Professor Engelhardt for histological analysis. *Tie2-ERT2/CRE^+^ Rosa26-DTR-STOP^fl/fl^* mice were newly crossed. For all other studies, we used male 8-10 weeks old C57BL/6 mice unless mentioned otherwise. Euthanizia of animals was performed by CO_2_ or by cervical dislocation.

### Senescence induction *in vitro*

To induce senescence by toxins, cells were plated subconfluent, and cultured in the presence of 1 μM palbociclib (PD033299, Pfizer Inc.), 100 nM doxorubicin (D1515, Sigma), or 10mU/ml bleomycin (B8416, Sigma) for 2 days, after which cells were washed, and replenished with complete culture medium. To induce senescence by iron, heme, or lysed RBCs, cells were plated subconfluent and cultured in 0,5% FBS containing RPMI medium (RPMI does not contain iron) for a day. Next day, cells were washed and replenished with 0.5% FBS containing RPMI medium which was supplemented with an iron solution 330 μM Fe(II)Sulfate (F8633, Sigma) and 330 μM Fe(III)Nitrate (F8508, Sigma), or lysed RBCs in 25-fold dilution (for the preparation, see next section), or with 10 μM hemin (H9039, Sigma Aldrich). After three days of culture, cells were washed and cultured in the presence of 10% FBS containing DMEM medium (contains free iron) for the time indicated. To induce senescence by irradiation, cells were plated subconfluent, and exposed to a dose of 20 Gy irradiation. To induce oncogene induced senescence, we used IMR90 cells stably transduced with a tamoxifen inducible Ras-G12V (kindly sent by Dr. Daniel Muñoz Espin) construct. Cells were cultured in the presence of 1 μM 4-hydroxy tamoxifen for 21 days.

### Preparation of lysed RBCs and considerations related to dosage of iron

Fresh blood was collected from mice in EDTA tubes. Blood was diluted in 1mM EDTA containing PBS, and the corpuscular fraction was isolated by three rounds of centrifugation at 600g. Corpuscular fraction was washed in EDTA-free PBS two times. PBS in the last centrifugation was removed, and replaced by a low volume of hypotonic 1x RBC lysis buffer (00-4333-57, eBioScience) (100 ul of RBC lysis buffer per 500 ul of corpuscular fraction). RBCs were vigorously vortexed, and passed through a gauge syringe multiple times, until a homogeneous, viscous red solution formed. This solution of lysed RBCs was resuspended in equal volumes of PBS, and supplemented to tissue culture in a 25- fold dilution to mimic micro-hemorrhagic or hemolytic insult.

We found that estimates in the literature for the iron content in RBCs from 1 ml of blood vary between 0,5-1 mg^2^, which would correspond to a molarity of 20-10 mM. To mimic this dosage of iron, our 1x iron solution was set to 16,5 mM. When comparing the effects of iron to those of lysed RBCs, we used the same dilutions of RBC and iron solutions. When delivering iron into the lungs, we delivered 30 μl of 16,5 mM iron solution to mimic a hemorrhagic insult caused by 30 μl of blood.

### Models of kidney fibrosis

For the folic acid induced kidney injury model, we used male C57BL/6 mice at 8-10 weeks of age were injected i.p. with a single high dose of folic acid (FA, 250mg/kg; F7876, Sigma) dissolved in 300mM sodium bicarbonate (NaHCO3; Sigma). For macroscopic analysis, both kidneys were taken. Dry kidney weight was normalized to the tibia length of the animals. For iron chelation, animals received daily intraperitoneal injections of 200 μl 1 mg/ml deferiprone (379409, Sigma) dissolved in water, control animals were injected with water only.

For the unilateral ureteral obstruction model we used male C57BL/6 mice at 8-10 weeks of age. Animals were anesthetized with isoflurane plus oxygen, and the left ureter was exposed through a flank incision, and the ureter ligated with 8–0 nylon, while in sham-operated mice the ureter was left undisturbed.

### Models of lung fibrosis

To induce pulmonary fibrosis 8- to 10-week-old male C57BL/6 wild-type mice were anesthetized by intraperitoneal injection with a mix containing ketamine (75 mg/kg) and medetomidine (1 mg/kg). Animals were placed on a Tilting WorkStand for rodents (EMC Hallowell) and intubated intratracheally with a 24GA catheter (BD Biosciences), and delivered 30 μl of bleomycin [1.5 U/kg of body weight] (Sigma, #15361), or 30 μl phosphate buffered saline (PBS), or 30 ul aqueous iron solution containing 8,3 mM Fe(II)Sulfate and 8,3 mM Fe(III)Nitrate (in total 16,5 mM iron). Mice receiving intratracheal iron developed within a day sepsis-like symptoms, with severe weight-loss, from which day recovered on the fourth day post-iron. We found that treatment with a daily dose of subcutaneous buprenorphine (0.1 mg/kg) considerably mitigated these symptoms, and the overall mortality in this model remained very low (<10%).

### Ischemic heart fibrosis

3-4 months old male C57BL/6J mice were used. Intraoperative analgesia was induced by treating mice with buprenorphine (0,1 mg/kg) 60 minutes prior to anesthesia. Anesthesia was induced with 5% isoflurane and it was maintained at 2% with mechanical ventilation following endotracheal intubation (2% isoflurane/97% oxygen, 130-140 stroke rate, stroke volume initially 5 ml/kg-increased to 7.5 ml/kg post-thoracotomy). Animals were placed on a heating table during the whole surgery and eyes were covered with ophthalmic gel (LACRYVISC 3 mg/g, Alcon) to protect the corneas. Neck hair and hair in the upper thorax was removed using a depilatory cream. An incision was made, followed by a left-sided thoracotomy at the fourth-intercostal space. The pericardium was removed and the heart left anterior descending artery was located between the pulmonary artery and the left auricle and was ligated using an 8-0 Prolene suture (PremiCron®- B Braun Surgical, S.A). The thoracic incision was closed in layers, using 6-0 Prolene sutures (PremiCron®- B Braun Surgical, S.A) to adapt the ribs and 4-0 Prolene sutures (PremiCron®- B Braun Surgical, S.A) to close the skin. A heating lamp was used to warm the animals until they fully recovered from the anesthesia. Analgesic treatment continued during the 3 following days after surgery by intraperitoneal injections of buprenorphine every 12h. Sham operated mice were used as control.

### Histochemical stainings

For hematoxylin and eosin (H&E) staining, paraffin-embedded tissue sections were dewaxed and stained with Hematoxylin eosin standard staining using a CoverStainer (Dako - Agilent). An Iron Stain Kit was used to identify iron pigments (AR15811-2, Artisan, Dako, Agilent) and a Masson’s Trichrome Stain Kit to identify collagen fibers and fibrin (AR17311-2, Artisan, Dako, Agilent). Stainings were performed in a Dako Autostainer Plus (Dako, Agilent) staining system following according to the manufacturer’s instructions. To enhance the iron detection, we adapted the enhanced Perl’s Prussian Blue staining protocol^3^ by adding a blocking step with 5% Normal Goat Serum (16210064, Life technology) with 2.5 % Bovine Serum Albumin (10735078001, Sigma) for 60 min followed with 30 min of Peroxidase Blocking Solution (S2023, Dako-Agilent) and a 30 min incubation with Liquid DAB+ Substrate Chromogen System (K3468, Dako-Agilent) on sections after that have been previously stained with the iron stain kit. For the Picrosirius Red Fast Green staining, tissue samples were incubated with the mordant Thiosemicarbazide 99% (TSC) (Sigma, T33405) for 10 min, washed in distilled water and therefore incubated with 0.1% Fast green (Sigma, FCF F7552) for 20 min and finally rinsed with 1% acetic acid (Sigma, 320099) for 1 min. In all cases samples were dehydrated and mounted with Mounting Medium, Toluene-Free (CS705, Dako, Agilent) using a Dako CoverStainer.

### Immunohistochemistry stainings

Immunohistochemistry was performed manually, in a Ventana discovery XT platform and in a Leica Bond RX platform . Primary antibodies were incubated on sections in a dilution and for the time as follows: p21 clone HUGO 291H/B5 (CNIO) RTU 60 min, p21 WAF1/Cip1 SX118 (Dako-Agilent, M7202) at 1:50 for 60min, NE (Abcam, ab68672) at 1:1000 for 120min, F4/80 D2S9R (CellSignalling, 70076) at 1:1000 for 60 min, alpha SMA [1A4] (Abcam, ab7817) at 1:750 for 120min, HMOX1 (Abcam, ab13243) at 1:1250 for 60 min and TER-119 (Stem Cell, 60033) at 1:5000 for 60min. Antigen retrieval for p21 was performed with Cell Conditioning 1 (CC1) buffer (Roche, 950-124). Secondary antibodies used were the OmniMap anti-Rat HRP (Roche, 760-4457) or OmniMap™ anti-Rb HRP (Roche, 760-4311). Blocking was done with Casein (Roche, 760-219). Antigen-antibody complexes were revealed with ChromoMap DAB Kit (Roche, 760-159). p21 WAF1/Cip1 SX118 were incubated with the rabbit to IgG1+IgG2a+IgG3 [M204-3] (Abcam, ab133469) at 1:500 for 32 min before adding the secondary. Prior to immunohistochemistry, sections were dewaxed and epitope retrieval was performed using Citrate buffer pH6 for α-SMA and Ter119 or Tris- EDTA buffer pH9 for HMOX1 in all cases for 20 min at 97°C using a PT Link (Dako, Agilent) or with ER2 buffer (AR9640, Leica) for 20 (F4/80) or 30 min (NE). Washings were performed using the Wash Solution AR (AR10211-2, Dako, Agilent) or with the BOND Wash Solution 10x (AR9590, Leica). Quenching of endogenous peroxidase was performed by 10 min of incubation with Peroxidase-Blocking Solution at RT (S2023, Dako, Agilent). Non-specific unions were blocked using 5% of goat normal serum (16210064, Life technology) mixed with 2.5 % BSA diluted in a wash buffer for 60 min at RT. Blocking of nonspecific endogenous mouse Ig staining was also performed using Mouse on mouse (M.O.M) Immunodetection Kit – (BMK-2202, Vector Laboratories). Secondary antibodies used were the BrightVision Poly-HRP-Anti Rabbit IgG Biotin-free, ready to use (DPVR-110 HRP, Immunologic), the HRP-Anti-Rat IgG (MP-7444, Vector) for 45 min or the Goat Anti-Mouse Immunoglobulins/HRP (Dako-Agilent, P0447) at 1:100 for 30 min. Antigen–antibody complexes were revealed with 3-3′- diaminobenzidine (K346811, Dako) or the DAB (Polymer) (Leica, RE7230-CE) with the same time exposure. Sections were counterstained with hematoxylin (CS700, Dako, Agilent) and mounted with Mounting Medium, Toluene-Free (CS705, Dako, Agilent) using a Dako CoverStainer. Specificity of staining was confirmed staining with the following isotype controls: rabbit IgG, polyclonal (Abcam, ab27478), the mouse IgG1, Kappa (NCG01) (Abcam, ab81032), mouse IgG2a kappa (eBM2a) (eBioscience™, 14-4724-82) and the rat IgG (R&D Systems, 6- 001-F).

### Image acquisition and quantification of histological sections

Brightfield images were acquired with a NanoZoomer-2.0 HT C9600 digital scanner (Hamamatsu) equipped with a 20X objective. All images were visualized with a gamma correction set at 1.8 in the image control panel of the NDP.view 2 U12388-01 software (Hamamatsu, Photonics, France). Image quantification was performed in the QuPath software^4^.

### SA-**β**-GAL staining

For senescence-associated ß-galactosidase staining cells were fixed in an SA-β-GAL fixation solution (PBS with 0.1M EGTA, 1M MgCl_2_ and 50% glutaraldehyde) for 15 minutes at room temperature and washed 2 times with PBS for a total 20 of minutes. Subsequently, a staining solution at pH6 was added (PBS with 1M MgCl2, 0.5M K_4_[Fe(CN)_6_] and 0.5M K_3_[Fe(CN)_6_] with 1 mg/ml X-Gal diluted in DMF (all Sigma)). The incubation took place overnight at 37 °C in a CO_2_ -free incubator.

### Measurement of free-hemoglobin in the serum

We collected serum samples from mice, and analyzed them for the levels of free hemoglobin by using the mouse hemoglobin ELISA kit (ab157715, Abcam), according to the manufacturer’s instructions.

### Ex vivo RBC hemolysis assay

Fresh blood was collected from mice in EDTA tubes. Blood was diluted in 1mM EDTA containing PBS, and the corpuscular fraction was isolated by three rounds of centrifugation at 600g. Corpuscular fraction was washed in EDTA-free PBS two times, and for the third time, resuspended in 4 volumes of PBS. 100 μl of RBC suspension was pipetted per well into a round bottom 96 well plate. Plate was centrifuged at 600 g, and the supernatant was replaced by a serial dilution of folic acid, by 0,05% Triton X-100 (T8787, Sigma) or by isotonic PBS. The plate was incubated for 6 hours at room temperature, centrifuged at 600 g to remove lysed RBCs, and sediment in the wells was resuspended in PBS.

### Bioinformatic analysis of lung biopsies - Rosa

We used gene set variation analysis (GSVA)^5^ to determine the relative expression scores of the GO_IRON_IRON_TRANSPORT gene ontology (GO: 0006826) in patients and controls from the two datasets of the lung tissue consortium (GSE47460). A kruskal-wallis test was used to compare the GSVA enrichment scores in controls vs. cases.

### qPCR

Total RNA was isolated using Trizol (Invitrogen) or the RNeasy Micro Kit (Quiagen) and cDNA was synthesized using the iScript cDNA synthesis kit (Bio-Rad). Quantitative realtime PCR was performed using the GoTaq SYBR Green qPCR Master Mix (Promega) and gene specific primers (**Extended Data Table 3.**). The relative abundance of transcripts was normalized to the expression of housekeeping genes using the 2^-ΔCT^ method. Relative expression values were calculated by normalizing gene expression to the average of the controls.

### Endothelial cell ablation in Tie2-ERT2/CRE^+^ Rosa26-DTR-STOP^fl/fl^ mice

Tamoxifen (Sigma) which was freshly dissolved in corn oil. For ablation of endothelial cells, mice were injected with tamoxifen (1 mg tamoxifen/20 g body weight, Sigma Aldrich), was administered to littermates by i.p. injection daily for 5 consecutive days.

### Western blotting

Cells were harvested after treatment and lysed in a buffer of 50 nM Tris-HCl pH 8, 1 mM EDTA, 150 mM NaCl, 1% NP40, 0,5% Triton X-100, 1% SDS with freshly supplemented with protease inhibitors (Roche#11873580001). Identical amounts of whole lysates (15 μg) together with a Chameleon™ Duo pre-stained protein ladder (LI-COR, #928-60000) were resolved in 4–12% Bis-Tris gels (NuPAGE Invitrogen, #NP0321BOX) and transferred to Amersham Protran 0.2 μm nitrocellulose membranes (GE Healthcare, #10600001). Blots were blocked with a LICOR blocking buffer, and incubated with the primary antibodies, namely anti-FTH1 (3998S, Cell Signaling) and anti-beta Actin (A5441, Sigma), on 4°C overnight, and subsequently incubated with the corresponding secondary anti-IgG antibodies anti-rabbit IRDye 680 CW (1:15,000, LI-COR, #926-68071), anti-rabbit IRDye 800 CW (1:15,000, LI-COR, #926-32211), and anti-goat IRDye 680 CW (1:10,000, LI-COR, #926-68074) for 1 h at room temperature. Blots were analyzed with an Odyssey Fc imaging system (LI-COR).

### Transmission electron microscopy

Cells were fixed at 4 °C with a mixture of 2% PFA and 2,5% glutaraldehyde in 0.1 M phosphate buffer (PB), then scrapped, pelleted, washed in PB, then postfixed with 1% osmium tetroxide and 0,8% potassium ferrocyanide. After that, samples were dehydratated with acetone and finally embedded in Spurr resin. Ultrathin sections were obtained using an Ultracut E (Reichert) and picked up on copper grids and observed with the JEM 1011 (JEOL, Japan) electron microscope, operating at 80kV. Micrographs were taken with a camera (Orius 1200A; Gatan, USA) using the DigitalMicrograph software package (Gatan, USA). Electron micrographs were processed using Adobe Photoshop CS6 (13.0.1) (Adobe Systems).

### Flow-cytometric analysis of lungs

Ex-vivo, lungs were intubated through the trachea with 2 ml dispase solution (5U/ml), and then agitated at 37°C. We chopped the lungs with fine scissors and transferred them into a solution of 1%BSA + 60units/ml DNase1 + 70units/ml Collagenase type 1 dissolved in PBS. To generate single-cell suspension, we used the GentleMACS system (Miltenyi) according to the manufacturer’s instructions. Red blood cells were removed in ammonium-chloride-potassium lysis buffer, and debris by using cell strainers. Cells were suspended in an ice-cold PBS with 1% FBS, before blocking with purified rat anti-mouse CD16/CD32 (BD). Surface antigens were stained with fluorescently conjugated antibodies: anti-CD45-PE, anti-CD45-APC, anti-CD11b-V450, anti-CD11b-PE/Cy7 and anti-Ly6G-PE all from BD, and F4/80-APC from Biolegend. Samples were acquired on a Gallios flow-cytometer (Beckman Coulter), acquired data was analyzed with a FlowJo software (Tree Star).

### Labile iron measurement

We measured labile iron by flow-cytometry using the FerroOrange Dye (F374, Dojindo) according to the manufacturer’s instructions.

### Total ROS measurement

Total ROS levels were measured by flow-cytometry using a total ROS detection kit (ENZ-51011, Enzo) according to the manufacturer’s instructions.

### Lysosomal mass measurement

Lysosomal mass was assessed by flow-cytometry using a LysoTracker dye (L12492, Invitrogen) according to the manufacturer’s instructions.

### Total iron

Total iron levels were measured by a colorimetric assay (MAK025-1KT, Sigma) according to the manufacturer’s instructions.

### Multiplex protein assays

For measuring protein levels in a multiplex format, lung tissues or cells were lysed in normal RIPA buffer, and protein concentrations were adjusted so that all samples contained the same amount of protein. Samples were shipped to an external commercial laboratory (EveTechnologies) and three types of routine discovery assays were performed on lungs tissues (MD44, MDCVD1, and MDMMP- C,O), and on type on senescence and non-senescent MEFs (MD44). Below we list the proteins for which these panels tested. Proteins not shown in the results were not changed significantly by the investigated insult.

**Mouse Cardiovascular Disease Panel 1 7-Plex Discovery Assay® Array (MDCVD1):** MMP-9, PAI-1 (Total), Pecam-1, sP-Selectin, sE-Selectin, sICAM-1, Thrombomodulin

**Mouse Cytokine/Chemokine 44-Plex Discovery Assay® Array (MD44):** Eotaxin, Erythropoietin, 6Ckine, Fractalkine, G-CSF, GM-CSF, IFNB1, IFNγ, IL-1α, IL-1β, IL-2, IL-3, IL-4, IL-5, IL-6, IL-7, IL-9, IL-10, IL-11, IL-12p40, IL-12p70, IL-13, IL-15, IL-16, IL-17, IL-20, IP-10, KC, LIF, LIX, MCP-1, MCP-5, M-CSF, MDC, MIG, MIP-1α, MIP-1β, MIP-2, MIP-3α, MIP-3B, RANTES, TARC, TIMP-1, TNFα, VEGF-A

**Mouse MMP 5-plex Discovery Assay® Array for Cell Culture and non-blood samples (MDMMP- C,O):** MMP-2, MMP-3, MMP-8, proMMP-9, MMP-12

### Diabetic nephropathy patients Perĺs

We performed Perl’s staining in 26 human biopsies with the diagnosis of diabetic nephropathy. Samples were collected between 2010 and 2016. The study was approved by the Bellvitge Hospital IRB. To selected samples having at least 10 glomeruli and patients with estimated glomerular filtration rate between 10 and 60 ml/min/1.73m^2^ at the time when the biopsy was performed. Iron staining was blindly evaluated by an investigator with experience in kidney pathology (JMC). Scoring was performed as follows. Interstitial fibrosis score: 0: absence; 1: 0-1/3 interstitial area; 2: 1/3-2/3 interstitial area; 3: 2/3-3/3 interstitial area. Glomerular sclerosis % of glomeruli with global sclerosis. Iron staining: 0: absence; 1: 0-10% tubuli; 2: >10% tubuli.

### Acute respiratory distress syndrome patients Perĺs

Autopsy study. An observational study was planned to evaluate the presence of iron in lung samples obtained from autopsies from patients who died with and without lung injury. The protocol was reviewed and approved by Comité de Ética de la Investigación Clínica del Principado de Asturias (ref 2020/151). Thirteen samples of paraffin-embedded lung tissue were obtained from autopsy material stored at the tissue bank at Hospital Universitario Central de Asturias (Oviedo, Spain). Clinical data was retrospectively collected. Acute respiratory distress syndrome (ARDS) was diagnosed in 7 cases, using the Kigali modification of Berlin criteria^6^ to allow the inclusion of patients without mechanical ventilation. Lung samples were taken from lower lobes. After deparaffinization, lung sections were stained with Perls Prussian blue and counterstained with eosin. Two researchers, blinded to clinical data, evaluated the extent of Perl’s stain from 0 (no visible staining) to 4 (presence of massive aggregates), and scores compared using a Wilcoxon’s test. In figures, the scale bar corresponds to 50 m.

### Magnetic resonance imaging of mice

MRI experiments were conducted on a 7.0-T BioSpec 70/30 horizontal animal scanner (Bruker BioSpin, Ettlingen, Germany) equipped with an actively shielded gradient system (400 mT/m, 12-cm inner diameter). Animals were placed in supine position in a Plexiglas holder with a nose cone for administration of anesthetic gasses (1.5% isoflurane in a mixture of 30% O_2_ and 70% CO) and were fixed using a tooth bar and adhesive tape. 3D localizer scans were used to ensure accurate position of the area of interest at the isocenter of the magnet. To estimate R2-map, an MSME (multi-slice mult- echo) sequence was acquired with 9 echo-times equally spaced from 11 to 99 ms, TR=3000 ms, acquisition matrix of 256x256, and voxel size of 0.12x0.12, slice thickness 0.8 mm. 10 slices were acquired resulting in a field of view of 30x30x8 mm³. MGE (multiple gradient echo) acquisition was performed to obtain R2* maps, with 10 echo-times equally spaced from 3.5 ms to 48.5 ms, TR=800 ms and 2 averages, acquisition matrix=256x256, voxel size of 0.12x0.12mm² and slice thickness 0.8 mm, 10 slices, FoV 30x30x8 mm². R2 and R2* maps were estimated using in-home scripts in Matlab (Mathworks Inc.). itkSNAP software (http://www.itksnap.org/) was used for segmentation and quantification: for each acquisition the kidneys’ cortex and medulla were manually delineated over the R2 map and overlaid on the R2* map to extract the average R2* values in these areas.

### Magnetic resonance imaging of renal allograft patients

#### Study design

Transversal study

#### Study intervention

MRI

#### Study population

Renal allograft recipients admitted to perform a kidney biopsy as standard of care clinical practice (per protocol or by clinical indication).

#### Inclusion criteria

- Written informed consent
- Age 18-80 year
- Kidney transplant recipient admitted to perform a kidney biopsy <u>Exclusion criteria:</u>
- Contraindications for MRI. Patients with pacemakers, defibrillators, other implanted electronic devices, metallic foreign body in the eye, insulin pumps.
- Pregnancy
- Absence of informed consent

#### Magnetic resonance imaging (MRI) procedure

All MRI studies were performed in a 1.5 Teslas scanner (Phillips Achieva d-Stream.Best, Netherlands) with a 32-channel phased-array surface coil. The MRI protocol included a sagittal T2 TSE weighted image covering the kidney allograft and perpendicular transverse slices acquired with Dixon Quant modified (mDixon Quant), an iron quantification sequence from de MRI vendor with four sets: water, fat, fat suppression and T2* images. Sequences were acquired with respiratory breath-hold and SENSE reconstruction. The mDixon Quant parameters were: six Echo Times with an initial TR of 5.3 and TE of 0.92, and the shortest repetition time. The matrix was 132x117 pixels and the field of view 400 x 350 mms. Flip angle of 5°. Bandwidth 2505.6 kHz. The scan mode was 3D, with 25 slices and slice thickness of 3.0 mms., gap 0. Number of signal averages (NSA) 1 and scan time duration 5,4 seconds. Three regions of interest (ROI) were manually delineated in the renal cortex taking care to avoid areas involved by susceptibility artifacts. They were placed in the anterior, posterolateral, and posteromedial parenchyma and copied in the four sets. T2* values were automatically calculated in the three ROIs and averaged to obtain a representative value for the kidney. The ROI main values were introduced in a web page calculator following de Wood Method to convert Hz/s in mol/g to quantify the iron burden.

#### Kidney allograft biopsy

The procedure was performed as per clinical practice. The histological sample was evaluated according to Banff 2017 score (Haas M et al, Am J Transplant 2018)

#### Estimation of sample size and statistical analysis

This is an exploratory study. We are going to include 20 kidney allograft recipients

Continuous variables with normal or symmetrical distribution were reported as mean and standard deviation above and below the mean. The categorical variables were described with frequencies and percentages. The differences between the two groups for continuous variables were analyzed using Student’s t test or the Wilcoxon rank-sum test, as appropriate. The differences between categorical variables were analyzed using the chi-squared distribution and Fisher’s exact test, as appropriate. Pearson’s and Spearman’s tests were utilized to assess the correlation between variables. All statistical tests were considered significant if the p value was <0.05 for two-tailed tests. The statistical analysis was performed with the Stat View SAS program.

#### Patients included

n=20.

#### Patients evaluated

n=13/20. Four patients were excluded because other MRI protocol/machine was used, 1 due to medical complication (dialysis initiation) and 2 because of withdrawal of consent.

